# Microbiome-Gut-Brain-Axis communication influences metabolic switch in the mosquito *Anopheles culicifacies*

**DOI:** 10.1101/774430

**Authors:** Tanwee Das De, Punita Sharma, Sanjay Tevatiya, Charu Chauhan, Seena Kumari, Deepak Singla, Vartika Srivastava, Jyoti Rani, Yasha Hasija, Kailash C Pandey, Mayur Kajla, Rajnikant Dixit

## Abstract

Periodic ingestion of a protein-rich blood meal by adult female mosquitoes causes a drastic metabolic change in their innate physiological status, which is referred to as ‘metabolic switch. Although the down-regulation of olfactory factors is key to restrain host-attraction, how the gut ‘metabolic switch’ modulates brain functions, and resilience physiological homeostasis remains unexplored. Here, we demonstrate that the protein-rich diet induces mitochondrial function and energy metabolism, possibly shifting the brain’s engagement to manage organismal homeostasis. A dynamic expression pattern of neuro-signaling and neuro-modulatory genes in both the brain and gut indicates an optimal brain-distant organ communication. Even after decapitation, significant modulation of the neuro-modulator receptor genes as well as quantitative estimation of neurotransmitters (NTs), together confer the gut’s ability to serve as a ‘second brain’. Finally, data on comparative metagenomic analysis and altered NTs dynamics of naïve and aseptic mosquitoes provide the initial evidence that gut-endosymbionts are key modulators for the synthesis of major neuroactive molecules. Conclusively, our data establish a new conceptual understanding of microbiome-gut-brain-axis communication in mosquitoes.

Graphical abstract

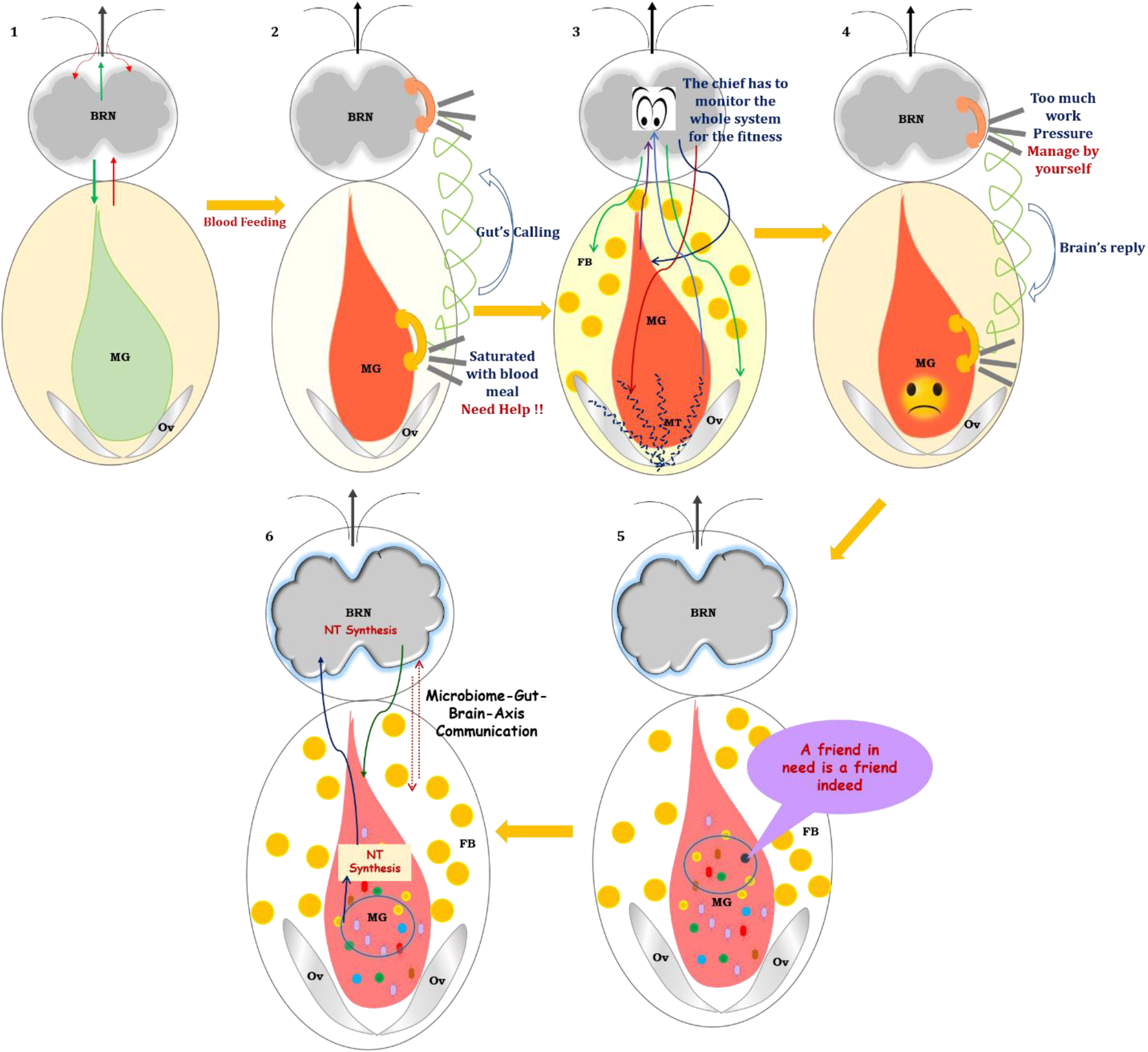

**Highlights:**

- Highly proteinaceous blood meal uptake causes gut ‘metabolic switch’ activity in mosquitoes.
- Gut’s calling shifts the brain’s administrative function from external communication to inter-organ management.
- ‘Gut’, as a ‘Second brain’ plays a crucial role in the maintenance of physiological homeostasis.
- Metabolic switch and proliferation of symbiotic bacteria establish microbiome-gut-brain axis communication in mosquitoes.

## Introduction

The brain is a privileged organ in shaping an animal’s behavior from lower to higher taxa by guiding and managing the diverse nature of external and internal stimuli. While each and every behavior of any organism is finely orchestrated by multiple organs, it is the brain that directs and exchange decision-making actions to regulate distinct organs functions. Unlike the human brain, which hosts billions of neurons, it is important to understand how insect’s especially blood-feeding mosquitoes having less than 1,00,000 neurons in their tiny brain, regulate similar functions such as finding a suitable source for sugar feeding and blood-feeding, searching for a mate-partner, and locating a proper oviposition site for egg-laying, etc. Decades of research have revealed that the molecular interaction of olfactory derived odorant-binding proteins (OBPs) and olfactory receptors (Ors) with the environmental chemical cues (external cues), play a central role in initializing these behaviors[1, 2]. However, successful completion of these behavioral activities is also influenced by the innate physiological status of the mosquitoes such as satiated/starved, mated/unmated, nutritional status, gravid, or not[3]. Thus how the miniature brain of mosquitoes harmonizes internal and external cues and affects decision-making abilities, is yet unknown.

Upon locating a suitable vertebrate host, a positive feeding decision stimulates the salivary glands of the adult female mosquito to facilitate rapid blood meal ingestion, and temporarily arrest olfactory actions until 30h of blood-feeding[1]. Although, a fully engorged female mosquito shows a dramatic suppression of host-seeking until the eggs are laid (~72 h post blood meal), however, the reactivation of the peripheral sensory system (30-40h post blood meal) is necessary to accompany complex oviposition site finding behavioral activities and gonotrophic cycle completion[4, 5]. A recent study by *Duvall et. al.* indicates that activation of Neuropeptide Y (NPY) signaling is essential for the suppression of host-seeking behavior for several days after blood feeding[4]. However, we have limited knowledge of how the mosquito’s brain regulates the binary behavioral switch responses (sugar to blood-feeding), and maintain organismal physiology in blood-fed females.

Fast engorgement of the mosquito’s gut with blood causes a drastic metabolic change in the innate physiological status from sugar to a protein-rich diet, resulting in the alteration of cellular fuel sources. This ‘metabolic switch’ drives multiple organs’ engagement to perform respective functions, such as osmoregulation by Malpighian tubules, progressive blood meal digestion by the gut, nutrient mobilization, and activation of vitellogenesis in the fat body, and ovary development for egg maturation[6–11]. The central nervous system should establish and maintain inter-organ communication to manage the sequential event of mosquitoes’ inner physiological activities. Several neuromodulators, such as neuropeptides, neurotransmitters, and neurohormones having a role in neuro-synaptic signal transmission and inter-organ communication, have been characterized in fruit fly[10, 12]. However, a similar correlation between the gut metabolic switch and brain function modulation in mosquitoes is limited to the *Aedes aegypti*, where brain secreted Insulin-like-peptide 3 is reported to play a significant role in the regulation of blood meal digestion and egg development [13]. Only a few recent genetic studies have suggested the key role of few neuropeptides e.g. Neuropeptide-Y, Short-neuropeptide F, and Allatostatin-A, and their receptors in the suppression of host-seeking and paternity enforcement in *Aedes aegypti* mosquitoes[4, 14, 15]. Furthermore, it is becoming increasingly evident in vertebrates that an enteric nervous system (vagus nerve), also referred to as ‘Second Brain’[16], not only mediates cross-talk between the gut and the brain but also establishes a bi-directional communication via gut-endosymbionts. This nexus of communication among the microbiota-gut-vagus-brain axis is crucial for maintaining metabolic homeostasis, mood, and perception[17–19]. Blood meal significantly modulates metabolic energy homeostasis in the mosquito gut, but how the gut’s nutrient-sensing mechanism influences brain function remains unknown[20]. Although blood-meal-induced gut-flora proliferation has been well demonstrated in mosquitoes[21], their neuromodulatory functions remain elusive.

Using a comprehensive RNA-Seq analysis of mosquito brain, coupled with extensive transcriptional profiling of neuro-modulators, comparative metagenomics analysis, and LC/MS-based quantitative estimation of neurotransmitters, here we demonstrated that fast blood meal engorgement and gut-metabolic switching (i) boost the brain’s energy-metabolism, which may likely influence organismal homeostasis, and (ii) favor the rapid establishment of a bidirectional microbiome-gut-brain axis communication, where the gut may also serve as a secondary brain in the blood-fed mosquitoes. Our data suggest that this gut-brain-axis communication is crucial for maintaining optimal physiological homeostasis during blood meal digestion and egg development in *Anopheles culicifacies* mosquitoes, the dominant malaria vector in rural India. A strategy of impairing this communication could reveal an out-of-the-box technique to disrupt mosquito host-seeking and blood-feeding behavior.

## Material and methods

**Fig. S1 represents a technical overview of the current investigation**

### Mosquito rearing and maintenance

A cyclic colony of *An. culicifacies* mosquito, sibling species A was reared and maintained at 28±2°C temperature and relative humidity of 80% in the central insectary facility of the ICMR-National Institute of Malaria Research. For routine rearing, adult female mosquitoes were fed on the rabbit. All protocols for rearing and maintenance of the mosquito culture were approved by the ethical committee of the institute.

### RNA isolation and transcriptome sequencing analysis

For RNA-Seq analysis, the brain tissues were dissected from 0-1-day old, 30 min post-blood-fed, and 30 h post-blood-fed cold anesthetized *An. culicifacies* mosquitoes by decapitation of the heads followed by application of gentle pressure over the head to pull out the brain tissue from the head cuticle and were collected in Trizol reagent. Then, total RNA was extracted from the collected brain tissues (approximately 30 mosquitoes were pooled to form one single sample), and a double-stranded cDNA library for each set of naïve, 30min, and 30h post-blood-fed were prepared by a prior well-established PCR-based protocol [22]. For transcriptome sequencing, the Illumina MiSeq 2 × 150 paired-end library preparation protocol was followed. The bioinformatics data analysis pipeline is shown in Fig S1. Briefly, raw reads from each set were processed to remove the adaptors and low-quality bases (<20). A de-novo clustering was used to build the final contigs/transcripts dataset using CLC Genomics Workbench (V6.2) (31) with default parameters (contig length ≥ 200, Automatic word size: Yes, Perform Scaffolding: Yes, Mismatch cost: 2, Insertion cost: 3, Deletion cost: 3, length fraction: 0.5, Similarity fraction: 0.8). Finally, the assembled transcriptome was used for CDS prediction and annotation using transdecoder software and BLASTX at e-value 1e^−6^ respectively. For a comprehensive differential gene expression analysis, we used the same protocol as mentioned previously [1, 22]. Additionally, to identify the differentially expressed genes associated with certain biological and molecular processes, we performed gene-list enrichment analysis using the Kobas 3.0 web server. The unique appearance of certain pathways in different brain samples was screened depending on the p-value (<0.5).

### PCR-based gene expression analysis

To establish the concept of the metabolic switch and inter-organ communication in mosquitoes, we targeted *An. culicifacies* brain, midgut, Malpighian tubule, and ovary tissues. The respective tissues were dissected and collected from both naïve sugar-fed and blood-fed mosquitoes originated from the same cohort at different time points. At first, the tissues were collected from 5-6-day old 25-30 naïve sugar-fed adult female mosquitoes. Next, adult female mosquitoes from the same cohort were offered blood-meal by offering a live animal (rabbit), and the desired tissues were collected as per the technical design of the experiments. In general, the fully engorged females were separated and kept in a proper insectary condition and the tissues were collected at the selected time points of post-blood-meal (PBM) such as 5min PBM, 2h PBM, 8-10h PBM, 24-30h PBM, 48h PBM, and 72h PBM from 25-30 mosquitoes for tissue-specific detailed expression analysis of the respective genes. The different tissues were pooled accordingly in Trizol and total RNA was extracted, followed by cDNA preparation. Differential gene expression analysis was performed using the normal RT-PCR and agarose gel electrophoresis protocols. For relative gene expression analysis, SYBR Green qPCR master mix and Biorad CFX 96 Real-Time PCR machine was used. PCR cycle parameters involved an initial denaturation at 95°C for 5 min, 40 cycles of 10 s at 95°C, 15 s at 52°C, and 22 s at 72°C. Fluorescence readings were taken at 72°C after each cycle. The final steps of PCR at 95°C for 15 secs followed by 55°C for 15 secs, and again 95°C for 15 secs were completed before deriving a melting curve. Each experiment was performed in three independent biological replicates for a better evaluation of the relative expression. The actin or Rps7 gene was used as an internal control in all the experiments, and the relative quantification was analyzed by the 2^−ΔΔCt^ method[23], which was further statistically analyzed by applying the student ‘t’ test and two-way ANOVA. The detailed list of primer sequences used in the study is mentioned in Table S1.

### ROS determination assay of blood-fed mosquitoes brain

To unravel the origin of the oxidative stress response in the blood-fed brain, we performed reactive oxygen species (ROS) determination assay by incubating the brain tissue dissected from naïve and blood-fed mosquitoes with a 2 mM solution of the oxidant-sensitive fluorophores, CM-H2DCFDA [5-(and-6)-chloromethyl-29,79-dichloro-dichlorofluorescein diacetate, acetyl ester] (Sigma). After a 20-min incubation at room temperature in the dark, the brain tissues were washed thrice with PBS, and then transferred to a glass slide in a drop of PBS and checked the fluorescence intensity at wavelength 490 nm under a fluorescent microscope.

### Antibiotic treatment of mosquitoes

To establish the concept of microbiome-gut-brain-axis communication, we disrupted the gut-commensal bacteria through antibiotic treatment. For the removal of gut bacteria, the pupae emerged in a washed and aseptic mosquito cage made up of muslin cloth. The antibiotic diet was provided to the newly emerged mosquitoes for 4-5days by mixing 10% sucrose solution with 10 μg of penicillin-streptomycin/ml and 15 μg gentamicin sulfate in it. To avoid any contamination, the antibiotic regimen was changed daily. After 4-5days of antibiotic treatment, blood-meal was provided to mosquitoes through the rabbit by maintaining proper sterile conditions such as (i) removed the extra hairs of rabbit pinnate/ears for easy access to blood meal, (ii) wipe the body of the rabbit with 70% ethanol, (iii) wipe the rabbit cage with alcohol.

### Decapitation Experiment

To test mosquito’s gut ability to function as a second brain, we offered a blood meal to 5-6 days old naïve sugar-fed mosquitoes, and decapitated ~100 mosquitoes after one hour of blood-feeding. Next, the decapitated mosquitoes were securely kept back in the insectary for recovery. The head tissues were submerged in 1X PBS to avoid desiccation. As per the technical design, post decapitation, the percentage of mosquito’s survival was recorded at different time points until we observed 100% mortality (mosquitoes that vibrate/move their legs or other body parts are considered as live and non-movable mosquitoes with visible shrinkage of the body parts at the respective time points are considered as dead). The brain and the gut tissues of surviving mosquitoes were dissected and collected at different time points for further gene expression analysis.

### Sample processing and MS analysis for neurotransmitter quantification

For absolute quantification of neurotransmitters, mosquitoes were decapitated and brains pulled out from the head cuticle and quickly collected in an Eppendorf containing 50μl of 1% ascorbic acid and immediately freeze it. For each set, ~60-65 mosquito brains or guts were pooled in a single tube. All samples were stored at −80°C until further use. Each sample was extracted with 3X volume of extraction solvent. Samples were vortexed and refrigerated for 10-15 minutes at 4°C. Samples were then subjected to sonication in a bath-type ultra-sonicator in pulses (twice, for 1 min each). Samples were then centrifuged at 14500 rpm for 5 mins at 4°C. The supernatants were separated and dried under a vacuum. Dried samples were spiked with internal standards (ISTDs) and derivatized, cleaned up, and prepared for LC-MS injections as per the protocol described earlier [24].

Briefly, Standards (STDs) were spiked in 200μl of extraction solvent (acidic acetone (0.1% FA) containing 0.5mM ascorbic acid) and dried under vacuum. ISTDs were spiked to dried STDs, followed by the addition of 80μL borate buffer (200mM, pH 8.8) containing 1mM ascorbic acid. To the above mixture 10μl of 0.1 N NaOH was added, followed by the addition of AQC (from 1 mg mL−1 stock). Samples were incubated at 55°C for 10 min. The reaction was stopped by the addition of 500μL of acidic water (0.1% FA). The derivatized standards were cleaned-up using the RP-SPE cartridges using the previously optimized protocol [25][24]: activation with methanol, equilibration with water (0.1% FA), loading of samples, washing (twice) with water (0.1% FA), and elution with acetonitrile: methanol (80:20) containing 2% FA. The eluate was dried under vacuum and reconstituted in 50μL of 0.5% acetonitrile. 10μL of reconstituted standards were injected for UHPLC-MS/SRM analysis.

Data were acquired on a TSQ Vantage (triple stage quadrupole) mass spectrometer (Thermo Fisher Scientific, San Jose, CA, USA) coupled with an Agilent 1290 Infinity series UHPLC system (Agilent Technologies India Pvt. Ltd.). The UHPLC system was equipped with a column oven (set at 40 °C) and a thermo-controller for maintaining the auto-sampler at 10 °C. A C-18 column (2.1 × 100 mm, 1.8μm, Agilent, Inc.) was used to perform the separation. The mobile phase solvent A was 10mM ammonium acetate in water containing 0.1% formic acid, and solvent B was acetonitrile containing 0.1% formic acid. The gradient was optimized to get maximum separation (2% B at 0 min, 2% B at 3 min, 20% B at 20 min, 35% B at 25 min, 80% B at 25–27 min, 2% B at 27–35 min) at a flow rate of 200μL min−1. The operating conditions were as follows: ionization mode: positive; spray voltage: 3700 V; capillary temperature: 270°C; source temperature: 80°C; sheath gas: 30 (arbitrary units); auxiliary gas: 10 (arbitrary units); collision gas: argon. Parent and product masses, S-lens voltages, and collision energies were used as per the previously optimized method [24, 25].

### Metagenomics analysis & microbiome profiling

For the metagenomics study, we collected gut from 3–4 days old sugar-fed adult female mosquitoes (n=50). While for blood-fed mosquito gut samples, 3-4 days old adult female mosquitoes from the same cohort were provided blood-meal by offering a live animal (rabbit), and midguts were collected after 24-30 hrs of blood-feeding. Before dissection, the mosquitoes were surface sterilized with 70% ethanol for 1 min in the highly sterilized condition of the Laminar airflow, and dissected tissues were collected in 1X Saline Tris-EDTA (100 mM NaCl/10 mM Tris-HCl, pH 8.0/1 mM EDTA, pH 8.0) buffer. At least 50 whole guts either from naïve sugar-fed or blood-fed mosquitoes, originating from the same cohort were collected into the minimal volume (20ul) of sterile ice-cold 1X STE and whole DNA was extracted as described earlier[26]. The quality of extracted gDNA was checked by loading the 5μl aliquot on 1 % agarose gel run at 110 V for 30 min. 1 μl of each sample was loaded in NanoDrop 8000 for determining the A_260/280_ ratio. The DNA was quantified using the Qubit dsDNA BR Assay kit (Thermo Fisher Scientific Inc.). 1 μl of each sample was used for determining concentration using Qubit^®^ 2.0 Fluorometer. For the preparation of amplicon libraries, V3-V4 hyper-variable region primers were used according to the library preparation protocol for the 16S Metagenomic Sequencing. The library for the sequenced fragments was obtained as per the standard Illumina protocol. QIIME software was used for maximum-likelihood phylogeny inference[27], generating OTUs for taxonomic identification and diversity estimation was performed using Megan software[28] (Table S3, Fig. S2). To validate the metagenomics data, the abundancy of the selected bacterial species was profiled through Real-Time PCR as described earlier[26].

## Results

### Hypothesis

Several independent studies highlighted the impact of age, sex, and circadian rhythm on the olfactory derived host-seeking behavioral properties of different mosquito species^26–28^. However, a holistic understanding of how a coordinated action of the neuro-olfactory system influences the host-seeking and blood-feeding behavioral properties remains unresolved^3^. Since the neuro-system is a highly sensitive and versatile center of chemical information exchange, we hypothesize that a minor change in the innate physiological status may have a strong impact on the mosquito’s everyday life. Importantly, after blood meal ingestion drastic changes in the innate physiological status and metabolic machinery i.e. “gut metabolic switch”, may engage multiple organs, to manage the events of the gonotrophic cycle. Therefore, it is very much plausible to propose that fast engorgement of mosquito gut with blood meal may shift mosquitoes’ brain functions from external communication to inter-organ management, such as (a) initiation of diuresis; b) finding a resting site for digestion of blood meal in the midgut; (c) distribution of amino acids, generated through the degradation of protein-rich blood meal; (d) active engagement of the fat body and ovary for egg maturation and life cycle maintenance (Fig. 1). We recently demonstrated that both mating and circadian have an important role in driving olfactory guided pre-blood-meal-associated behavioral properties even ‘Without Host Exposure’ in the aging adult female mosquito *An. culicifacies*[1, 29, 30]. Here, we aimed to decode and establish a possible molecular correlation between brain and gut-metabolic switch in *An. culicifacies*. First, to unravel the brain’s molecular complexity in response to blood feeding, we followed a similar RNA-Seq strategy as planned for the olfactory system[1], as detailed in Fig.1.

**Fig. 1:**
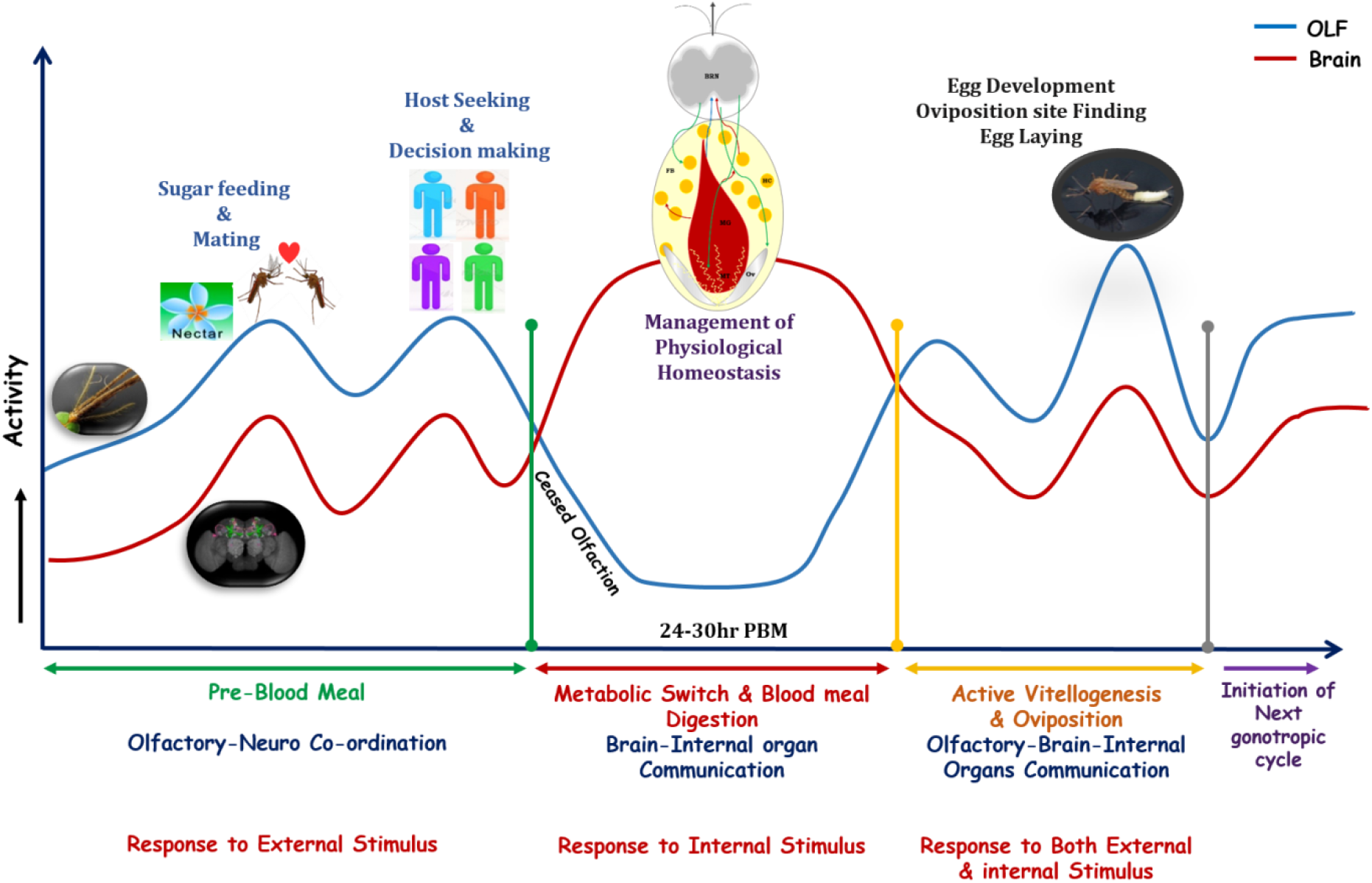
A proposed working hypothesis to establish the correlation between the gut metabolic switch and brain functions in adult female mosquitoes. The behavior of any organism is a very complex event that needs tight coordination between the sensory and neuronal systems. After emergence from pupae, the dynamic changes in the neuro-olfactory system coordinate and regulate different behavioral activities such as mating, sugar feeding, and vertebrate host-seeking, etc. These pre-blood meal-associated behaviors are guided by external stimuli, followed by neuronal decision making. Once the female mosquitoes take a blood meal, their olfactory responses are temporarily arrested to minimize brain and environmental (external) communication. But blood-feeding causes a global change in physiological homeostasis and drives multiple tissues (midgut, Malpighian tubule, ovary, and fat body) engagement to manage the systemic equilibrium. Here, we hypothesize that an ‘*internal stimulus*’ of gut-metabolic-switch may modulate brain functions to ensure optimal inter-organ communication at least for the first 30h until blood meal digestion is completed in the gut. However, after 30-40h of blood-feeding reactivation of the olfactory system, restores olfactory-neuro co-ordination to perform the next level of behavioral activities, such as oviposition and initiation of the second gonotrophic cycle. Blue and red lines indicate the possible functional patterns of the olfactory system (OLF) and the brain, respectively.

### Blood meal ingestion boosts the brain’s energy metabolism

A comparative analysis of naïve sugar-fed, 30min, and 30hpostblood-fed mosquito’s brain RNA-Seq data showed a gradual suppression of brain-specific transcript abundance (Fig. 2a). Surprisingly, we also observed an exceptional enrichment of oxidation-reduction process associated transcripts in response to blood-feeding (Fig. 2b) (Table S2a, S2b), though, we failed to detect any signal of oxidative stress in a 2mM solution of the oxidant-sensitive fluorophores, CM-H2DCFDA [5-(and-6)-chloromethyl-29,79-dichloro-dihydrofluorescein diacetate, acetyl ester] (data not are shown). While an in-depth analysis of oxidation-reduction category transcripts showed enrichment of several mitochondrial activity proteins such as 2-oxoglutarate dehydrogenase, NADH dehydrogenase, glutathione peroxidase, etc. A comparative metabolic pathway prediction analysis further confirmed the exclusive induction of several unique pathways linked to (a) energy metabolism, (b) neurotransmitter synthesis, and (c) neurite outgrowth and synaptic transmission (Fig. 2c). Together these data indicated that blood meal associated gut metabolic switch may trigger a “hyper energy” state and neuro-modulatory factors in the mosquito brain.

**Fig. 2:**
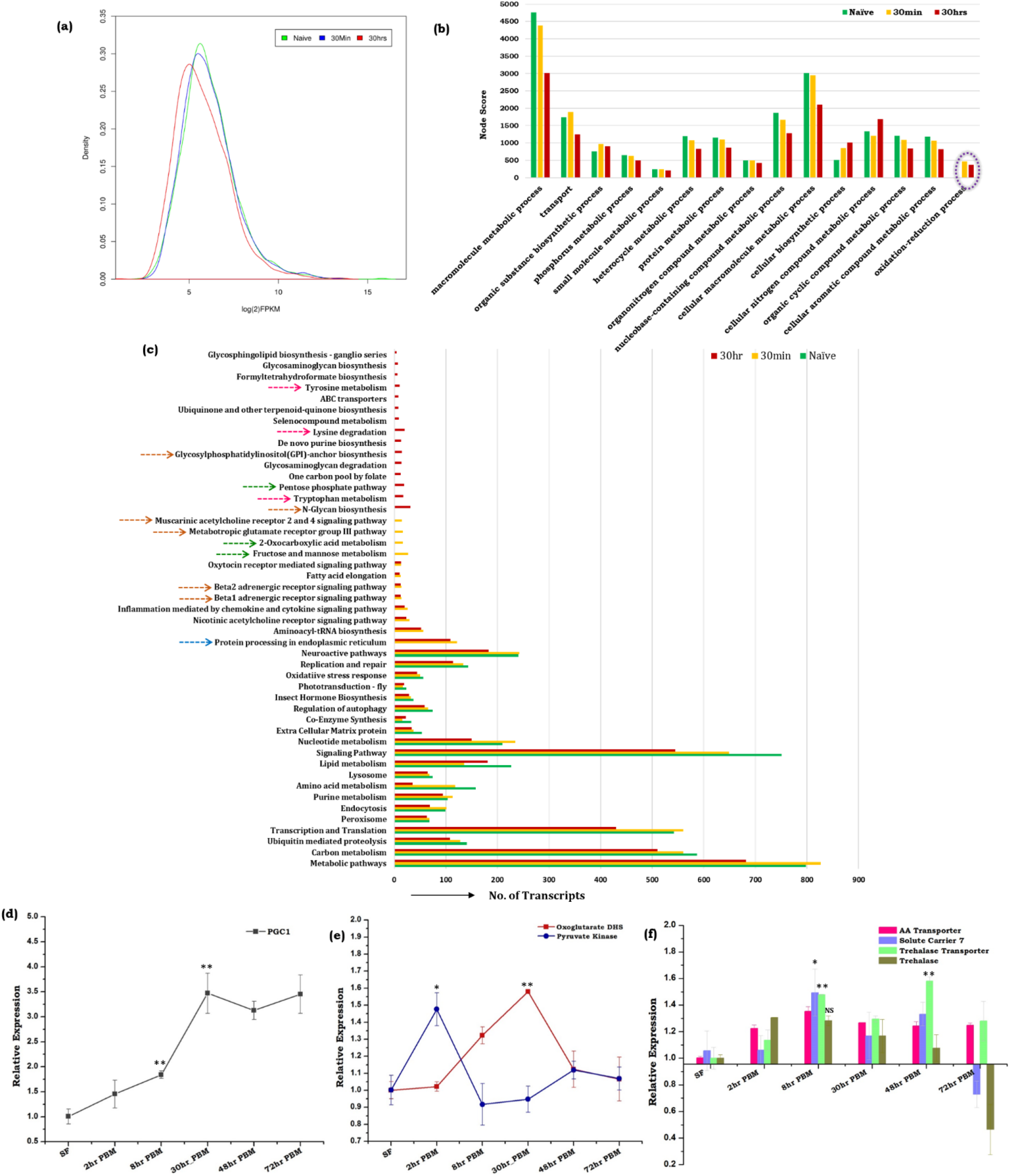
Blood meal causes notable changes in the molecular architecture of the brain tissue. (a) Comparison of the read density map of the naïve, 30min, and 30h post blood meal (PBM) transcriptomic data of brain tissue (n=25); (b) Functional annotation and molecular cataloging of brain transcriptom**e**(Biological Process/Level4/Node score). Purple circle highlighted the unique category of genes that appeared in the brain tissue after blood meal intake; (c) KOBAS 3.0 software mediated gene list enrichment and comparative pathway analysis of naïve and blood-fed brain tissues. Green arrow links to energy metabolic pathways, the pink arrow links to neurotransmitter synthesis pathway, and brown arrow indicate neurite out-growth and synaptic transmission; (d) Relative expression profiling of PGC-1 gene in the brain of naïve and blood-fed mosquitoes*(n* = *25, N* = *3);* (e) Transcriptional profiling of transcripts related to energy metabolism in the brain tissue of naïve and blood-fed mosquitoes at different time points; (f) Comparative transcriptional response of amino acid transporters and trehalose transporter along with trehalase enzyme in the brain tissue after the metabolic switch *(n* = *25*, *N* = *3)*. Statistically significant variation in the expression of the respective genes was tested by the *t-test* and compared with the sugar-fed control brain. (n = number of mosquitoes from, which the respective tissue was dissected and pooled for each independent experiment; N = number of biological replicates)

To verify the above presumption, we profiled and compared the expression pattern of the PGC-1 gene (Peroxisome proliferator-activated receptor gamma coactivator 1-alpha), an important transcriptional co-activator that regulates genes involved in energy metabolism[31–33]. A persistent elevation of PGC-1 (P ≤ 0.009 at 8h PBM, P ≤ 0.007 at 30h PBM), and a parallel enrichment of glycolysis and TCA cycle gene pyruvate kinase (P ≤ 0.0176) and oxoglutarate dehydrogenase (P ≤ 0.0019) respectively, indicated an enhanced mitochondrial activity in the brain of blood-fed mosquitoes (Fig. 2d, e). Next, we tested whether the amino acids generated through blood meal digestion or trehalose, a non-reducing disaccharide, acts as raw material for the brain’s energy metabolism. Although trehalose serves as a primary energy source in the insects’ brain[34, 35], we observed a sequential increment in the amino acid (P ≤ 0.0515) as well as trehalose (P ≤ 0.0071) transporter genes in the blood-fed brain (Fig. 2f). Together, these data indicate that both amino acids and trehalose may synergistically communicate the nutritional signal to the brain for active management of multi-organ communication[13, 36].

### Spatial and temporal modulations of neuro-signaling regulate metabolic switch-associated physiological activities

In naive sugar-fed mosquitoes, olfactory guided neuro-signaling and the brain’s energy consumption is optimal to drive external stimuli associated with routine behavioral events like flight, mating, and host-seeking. Recently, we have demonstrated that prior host-exposure, sex, and circadian have a significant impact on the olfactory responses in the aging adult-female mosquito *An. culicifacies*[1, 29, 30], Likewise, to test and evaluate the possible correlation of age with neuro-regulation, we monitored the expression profile of at least 14 neuro-modulatory genes in aging naïve adult female mosquito’s brain (Fig. S3). A limited modulation of expression suggested that host exposure may have an important role in cognitive learning and blood-feeding behavioral adaptation in mosquitoes, though further studies are needed to clarify and establish the correlation. However, surprisingly, after blood feeding an increase in the brain’s energy consumption, prompted us to test the functional correlation of the brain with gut metabolic switch activities. Here, we hypothesize that blood meal uptake may temporarily pause the external communication, and increased energy state possibly may favor the shifting of the brain’s engagement for the maintenance of organismal homeostasis. Thus, we identified and shortlisted transcripts encoding proteins, likely involved in the key events of the synaptic signal transmission process, i.e., crucial for the brain’s functioning. Thereafter, we evaluated the blood-meal-associated transcriptional response of selected transcripts regulating either receptor-mediated neuronal or cellular signaling processes during synaptic transmission.

Surprisingly, we observed a limited change in the expression of neurotransmitters and biogenic amine receptor genes such as serotonin receptor, dopamine receptor, octopamine receptor, and GABA receptor, etc. in response to blood-meal (Fig. 3a). While, on the contrary, cellular signal transduction proteins such as cGMP protein kinase, phospholipase C, GABA gated chloride channel, and serine-threonine protein kinase, exhibited a significant modulation in response to metabolic switch (Fig. 3b). Together, these findings support the idea that a rapid blood meal ingestion may drive brain engagement to manage metabolic switch-associated activities and distant organs’ function.

**Fig. 3:**
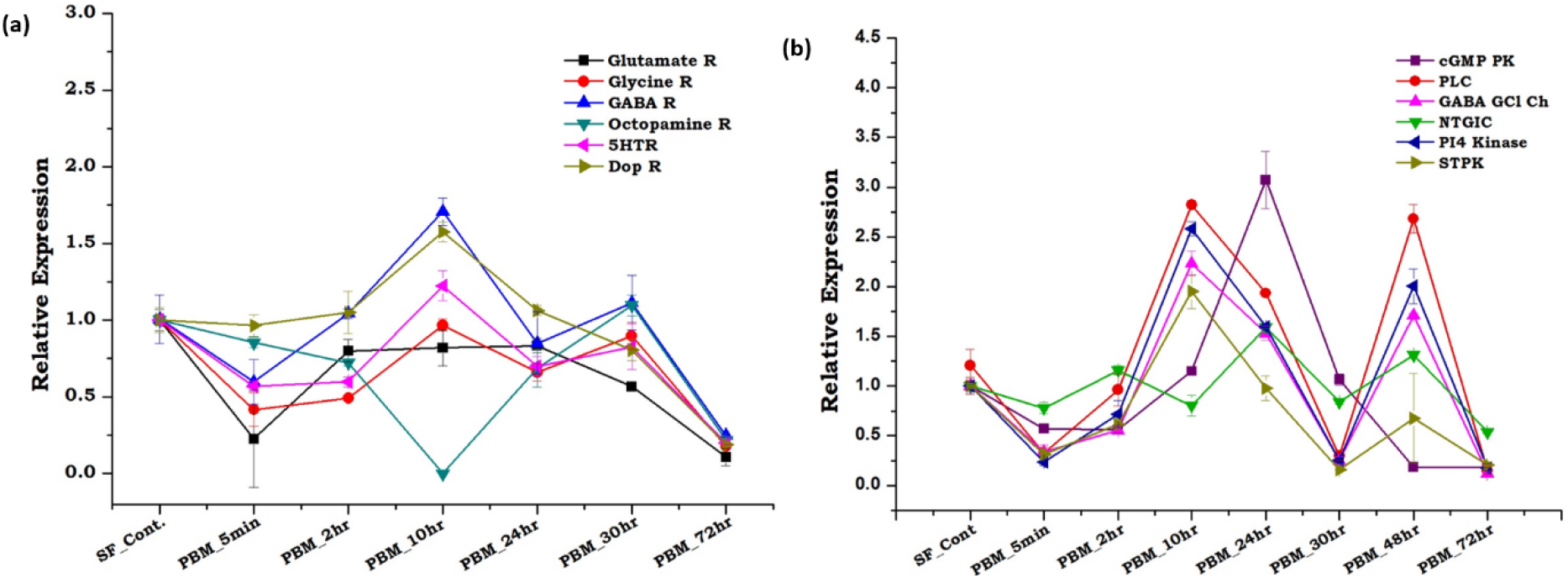
Metabolic switch influences neuro-signaling modulation. (a) Transcriptional response of neurotransmitter receptor genes as per the designed blood meal time-series experiment. Glutamate R: Glutamate Receptor; Glycine R: Glycine Receptor; GABA R: Gamma-Aminobutyric Acid Receptor; Octopamine R: Octopamine Receptor; 5HTR: Serotonin Receptor; Dop R: Dopamine Receptor. Statistical analysis using two-way ANOVA has implied at 0.05 level the expression pattern of the respective genes was not statistically significant at p ≤ 0.2 at different time points after blood feeding *(n* = *25, N* = *4);* (b) Relative expression profiling of the genes involved in signal transduction molecules according to the detailed blood meal time-series experiment. cGMP PK: Cyclic GMP Protein Kinase; PLC: Phospholipase C; GABA GClCh: GABA Gated Chloride Channel; NTGIC: Neurotransmitter Gated Ion Channel; PI4 Kinase: Phosphatidyl-inositol-4-Kinase; STPK: Serine Threonine Protein Kinase. Statistical analysis using two-way ANOVA stated that the expression change of the respective genes is statistically significant p ≤ 0.005 *(n* = *25, N* = *4)*. (n = number of mosquitoes from which the respective tissue was dissected and pooled for each independent experiment; N = number of biological replicates)

### Innate physiological status differentially modulates tissue-specific neuromodulators/receptors transcripts expression

To further validate and correlate brain-inter-organ communication, we monitored the temporal and spatial expression of at least 21 key genes (Table 1) having the blood-meal-associated function in their targeted tissue such as midgut (MG), ovary (Ov), and Malpighian tubules (MT). Notably, we observed a significant upregulation of ILP3(p<0.0002) and also time-dependent modulation of other neuropeptides (Neuropeptide Y, Leukokinin) and neuro-hormones (OEH, DH44, and ARMAA) in the blood-fed mosquitoes brain (Fig. 4a, b, c). We interpreted that a gradual induction of ILP3 synthesis and OEH secretion from the brain’s neurosecretory cells may activate the ovaries for the synthesis of ecdysteroids to initiate the vitellogenesis process [37–39]. A transient increase in NRY immediately after blood-feeding may be due to gut distension, but a significant increase (P<0.005) after 24h and 72h may cause suppression of host-seeking, a mechanism recently reported in *Aedes aegypti*[4, 40].

**Table 1:**
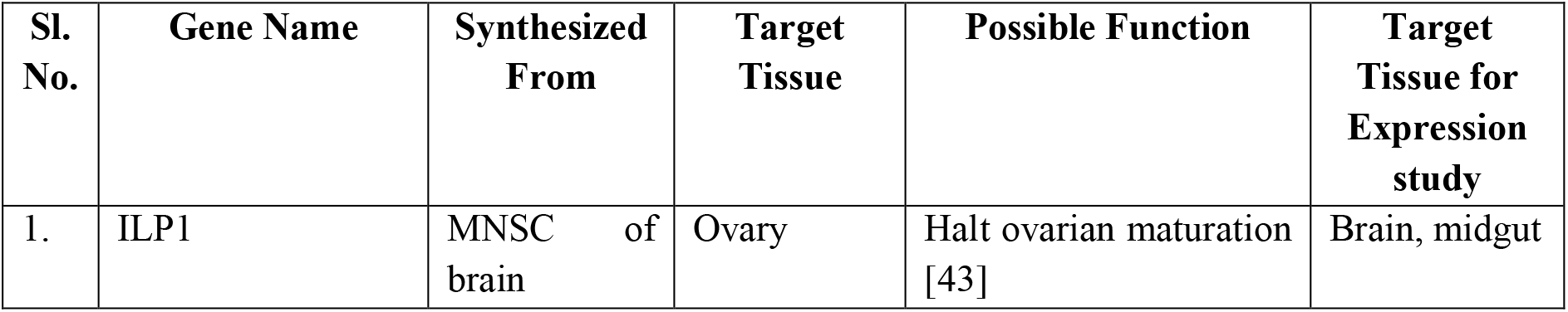
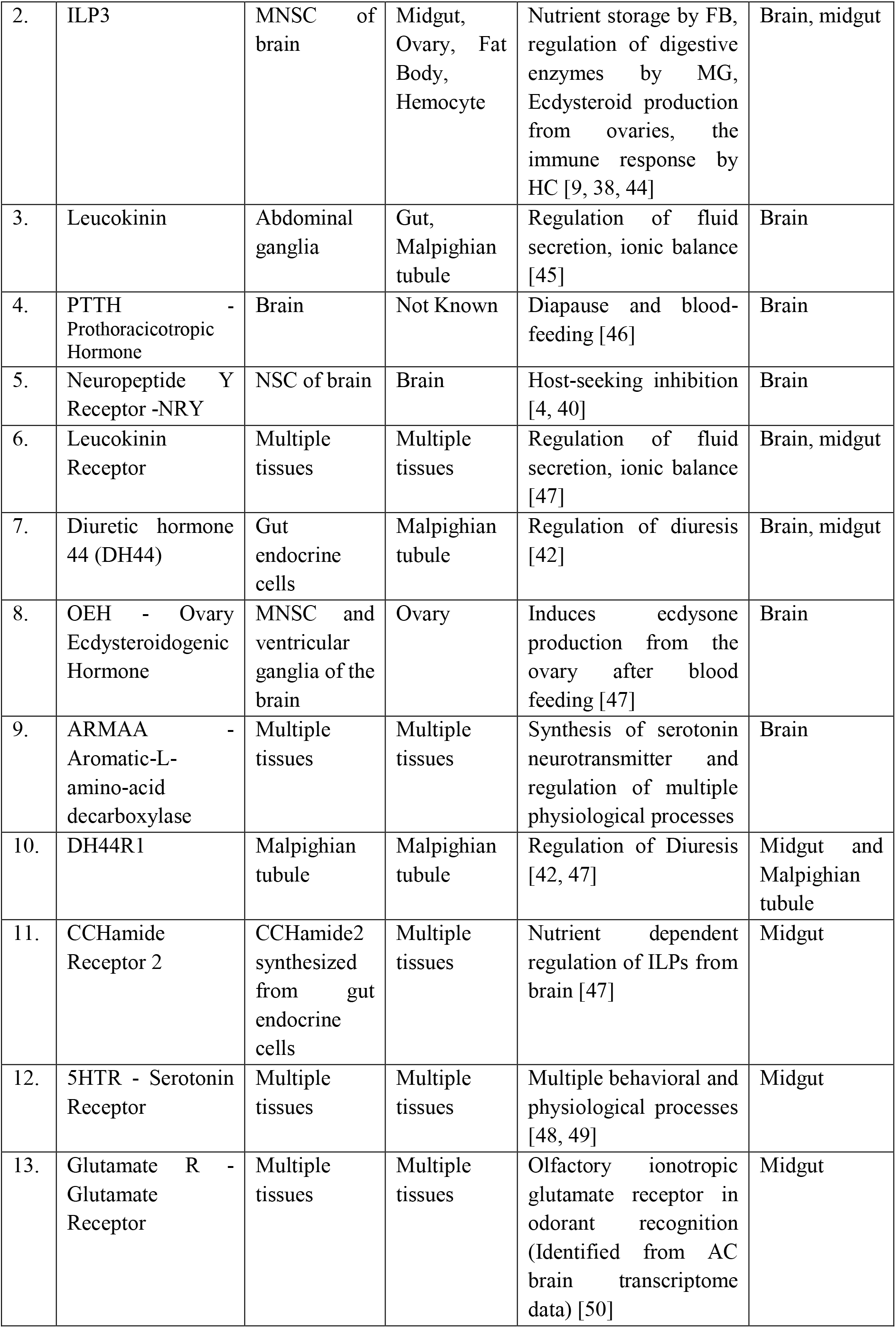
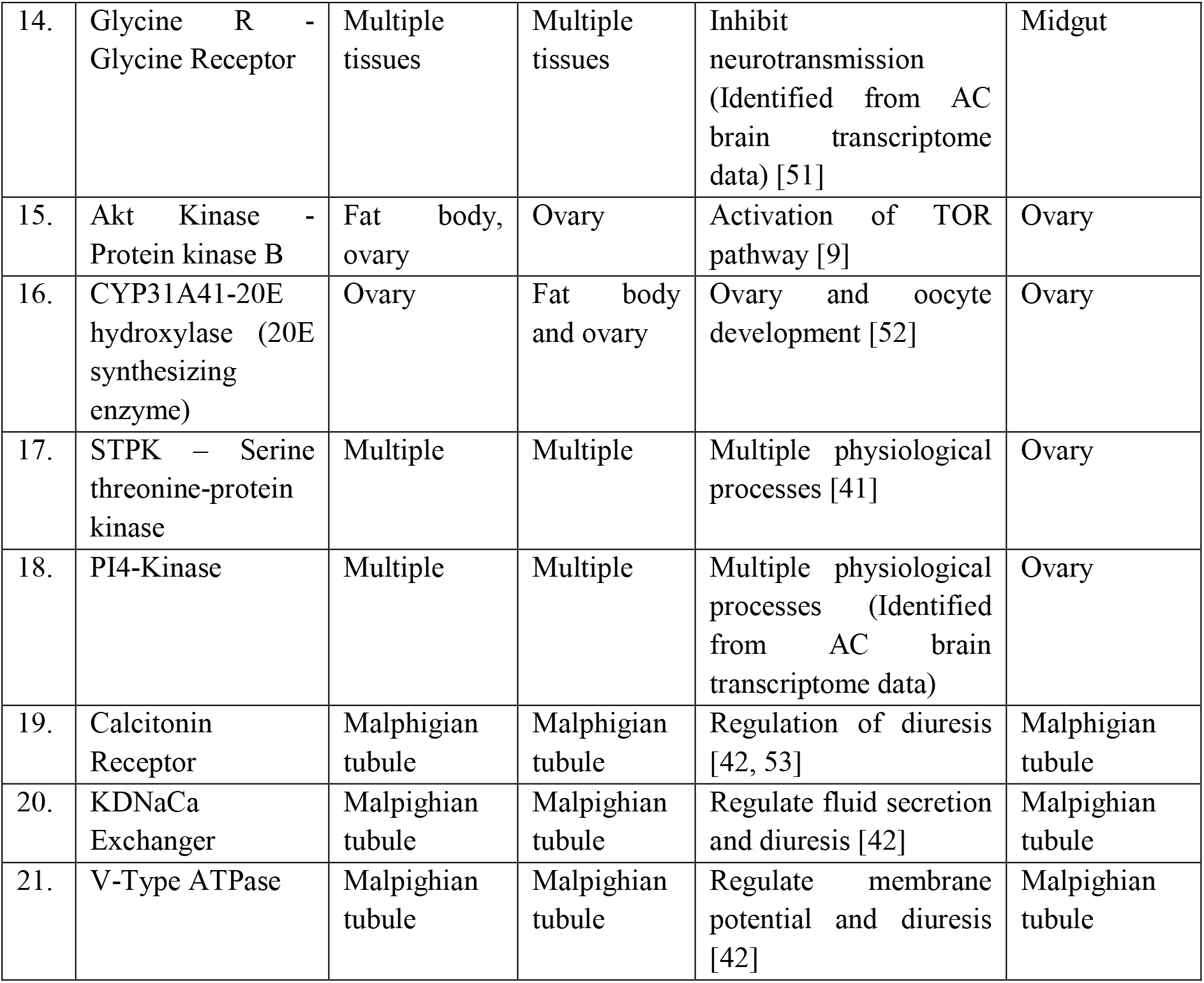
Details of the selected transcripts used to understand inter-organ communication during metabolic switch events.

| Sl. No. | Gene Name | Synthesized From | Target Tissue | Possible Function | Target Tissue for Expression study |
| --- | --- | --- | --- | --- | --- |
| 1. | ILP1 | MNSC of brain | Ovary | Halt ovarian maturation [43] | Brain, midgut |
| 2. | ILP3 | MNSC of brain | Midgut, Ovary, Fat Body, Hemocyte | Nutrient storage by FB, regulation of digestive enzymes by MG, Ecdysteroid production from ovaries, the immune response by HC [9, 38, 44] | Brain, midgut |
| 3. | Leucokinin | Abdominal ganglia | Gut, Malpighian tubule | Regulation of fluid secretion, ionic balance [45] | Brain |
| 4. | PTTH - Prothoracicotropic Hormone | Brain | Not Known | Diapause and blood-feeding [46] | Brain |
| 5. | Neuropeptide Y Receptor -NRY | NSC of brain | Brain | Host-seeking inhibition [4, 40] | Brain |
| 6. | Leucokinin Receptor | Multiple tissues | Multiple tissues | Regulation of fluid secretion, ionic balance [47] | Brain, midgut |
| 7. | Diuretic hormone 44 (DH44) | Gut endocrine cells | Malpighian tubule | Regulation of diuresis [42] | Brain, midgut |
| 8. | OEH - Ovary Ecdysteroidogenic Hormone | MNSC and ventricular ganglia of the brain | Ovary | Induces ecdysone production from the ovary after blood feeding [47] | Brain |
| 9. | ARMAA - Aromatic-L-amino-acid decarboxylase | Multiple tissues | Multiple tissues | Synthesis of serotonin neurotransmitter and regulation of multiple physiological processes | Brain |
| 10. | DH44R1 | Malpighian tubule | Malpighian tubule | Regulation of Diuresis [42, 47] | Midgut and Malpighian tubule |
| 11. | CCHamide Receptor 2 | CCHamide2 synthesized from gut endocrine cells | Multiple tissues | Nutrient dependent regulation of ILPs from brain [47] | Midgut |
| 12. | 5HTR - Serotonin Receptor | Multiple tissues | Multiple tissues | Multiple behavioral and physiological processes [48, 49] | Midgut |
| 13. | Glutamate R - Glutamate Receptor | Multiple tissues | Multiple tissues | Olfactory ionotropic glutamate receptor in odorant recognition (Identified from AC brain transcriptome data) [50] | Midgut |
| 14. | Glycine R - Glycine Receptor | Multiple tissues | Multiple tissues | Inhibit neurotransmission (Identified from AC brain transcriptome data) [51] | Midgut |
| 15. | Akt Kinase - Protein kinase B | Fat body, ovary | Ovary | Activation of TOR pathway [9] | Ovary |
| 16. | CYP31A41-20E hydroxylase (20E synthesizing enzyme) | Ovary | Fat body and ovary | Ovary and oocyte development [52] | Ovary |
| 17. | STPK – Serine threonine-protein kinase | Multiple | Multiple | Multiple physiological processes [41] | Ovary |
| 18. | PI4-Kinase | Multiple | Multiple | Multiple physiological processes (Identified from AC brain transcriptome data) | Ovary |
| 19. | Calcitonin Receptor | Malpighian tubule | Malpighian tubule | Regulation of diuresis [42, 53] | Malpighian tubule |
| 20. | KDNaCa Exchanger | Malpighian tubule | Malpighian tubule | Regulate fluid secretion and diuresis [42] | Malpighian tubule |
| 21. | V-Type ATPase | Malpighian tubule | Malpighian tubule | Regulate membrane potential and diuresis [42] | Malpighian tubule |

**Fig. 4:**
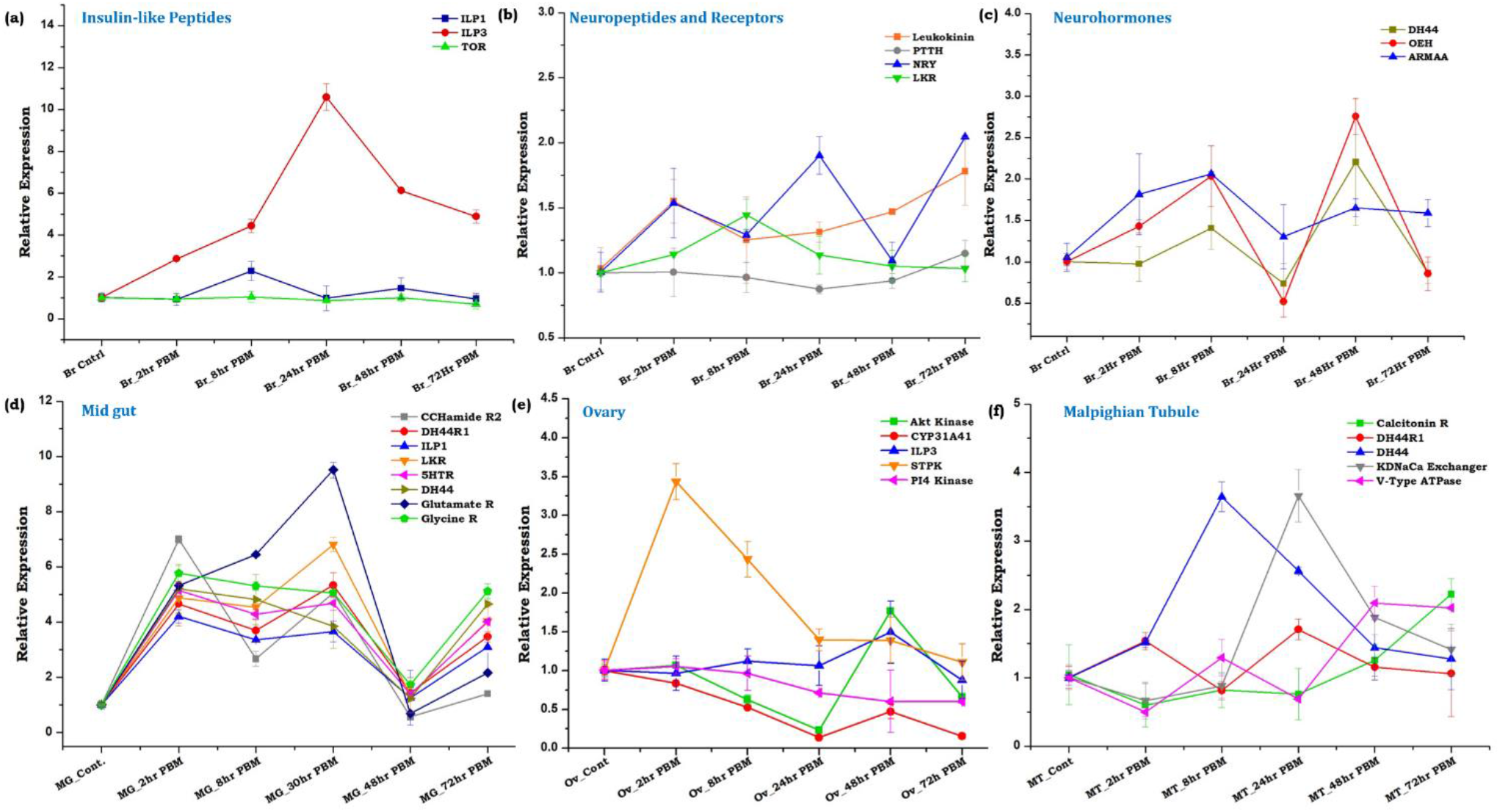
Metabolic switch modulates tissue-specific neuro-modulator transcripts expression. (a-c) Transcriptional expression profiling of Insulin-like-peptides, neuropeptides, neurohormones, and receptor genes in the brain tissue during the metabolic switch. Statistical analysis using two-way ANOVA and Tukey’s test implied that the expression change of the respective genes is statistically significant for insulin-like-peptides p ≤ 0.007; neuropeptides and receptors p ≤ 0.009, but for neuro-hormones, it was non-significant p ≤ 0.2 *(n* = *25, N* = *4);* (d) Relative expression profiling of a subset of neuromodulator genes in the midgut of naïve and blood-fed mosquitoes at the same time point described above. Statistical analysis using two-way ANOVA implied that the expression change of the respective genes is statistically significant p ≤ 0.005 *(n* = *12, N* = *4);* (e) Transcriptional profiling of genes involved in signal transduction during vitellogenesis in the ovary. Statistical analysis using two-way ANOVA and Tukey’s test indicated that the expression change of the respective genes was statistically significant at p ≤ 0.002 *(n* = *12, N* = *4);* (f) Relative gene expression analysis of diuresis-related genes in the Malpighian tubule of naïve and blood-fed mosquitoes. Statistical analysis using two-way ANOVA and Tukey’s test indicates that the expression change of the respective genes is non-significant at p ≤ 0.4 *(n* = *25, N* = *4)*. (n = number of mosquitoes from which the respective tissue was dissected and pooled for each independent experiment; N = number of biological replicates)

Next, we asked how the dynamic changes of the neuromodulators in the blood-fed brain influence distant organs responses, such as diuresis regulation by the Malpighian tubule, blood digestion process in the midgut, and oocyte maturation in the ovary. Transcriptional profiling of selected neuropeptide, neurotransmitter receptor transcripts (Table 1) indicated that blood meal triggers an immediate and prolonged (~48h PBF) impact on the expression of the gut-neuro transcript (Fig. 4d). Parallel observation of an early induction (2h PBF) of serine threonine-protein kinase (MAPK activated protein kinase) and late expression of Akt kinase (48h PBF) in the ovary suggested a controlled regulation of the nutritional signaling pathway favors the vitellogenesis process (Fig. 4e) [9, 41]. Likewise, observation of a unique pattern of diuretic hormone (8h PBF) and potassium dependent sodium-calcium exchanger gene (24h PBF) expression in the Malpighian tubule suggested an active diuresis process until 24h post blood meal (Fig. 4f) [42].

### Gut, the ‘*second brain*’ communicates the nutritional status through neurotransmitter synthesis

In vertebrates and also in the fruit flies, it is well evident that effective communication between the gut and brain has a paramount effect in shaping optimal health[16, 18], but a very limited knowledge exists in the mosquitoes[13]. Prolonged modulation of the neuromodulators expression in the blood-fed mosquitoes’ gut invigorates us to presume the existence of bi-directional gut-brain axis communication. An enriched expression pattern of neurotransmitter receptor genes, even after decapitation, reflected that the gut may also perform neuro-modulatory actions independently (Fig. S3). To further establish a proof-of-concept, we followed LC/MS-based absolute quantification of different neurotransmitters (NT) and compared their levels in the brain as well as in the gut of naïve and blood-fed mosquitoes.

Our data revealed that in naïve sugar-fed mosquitoes, although the brain serves as the primary source of NT synthesis, the midgut also synthesizes a substantial amount of NTs (Fig. 5a). However, blood-feeding causes a drastic shift in the NTs level in the midgut than in the brain (Fig. b, c). Notably, we observed an unpredictable increase in most NTs except glutamic acid, tyrosine, and tyramine in the gut (Fig. 5c). Whereas, the brain tissue showed a notable decrease in the majority of the NTs synthesis, except for histamine, tyrosine, and tryptophan (Fig. 5b). We also observed that tyrosine amino acid was exclusively induced in the brain after blood-feeding, but remained below the threshold level in the gut (Fig. 5 b, c). Although our data support previous studies that in addition to the brain, the gut also serves as a major source of multiple neurotransmitters in vertebrates and fruit fly[16, 54], the mechanism of nutrition-dependent NTs modulation remains unclear. Especially, in mosquitoes our understanding of the complex nature of blood meal digestion and gut-brain axis communication is obscure. Thus, our unusual observation of a thousand-fold increase in the levels of histidine, serine, aspartic acid, and tryptophan in the blood-fed mosquito’s gut emanated few key questions: 1) whether increased levels of amino acids in the gut during blood meal digestion may act as an NT? 2) Do blood-meal-induced proliferation of the gut microbiota has any effect on NT dynamics? 3) Do the gut endosymbionts of mosquitoes have any impact on gut-brain axis communication? (Fig. 5d).

**Fig. 5:**
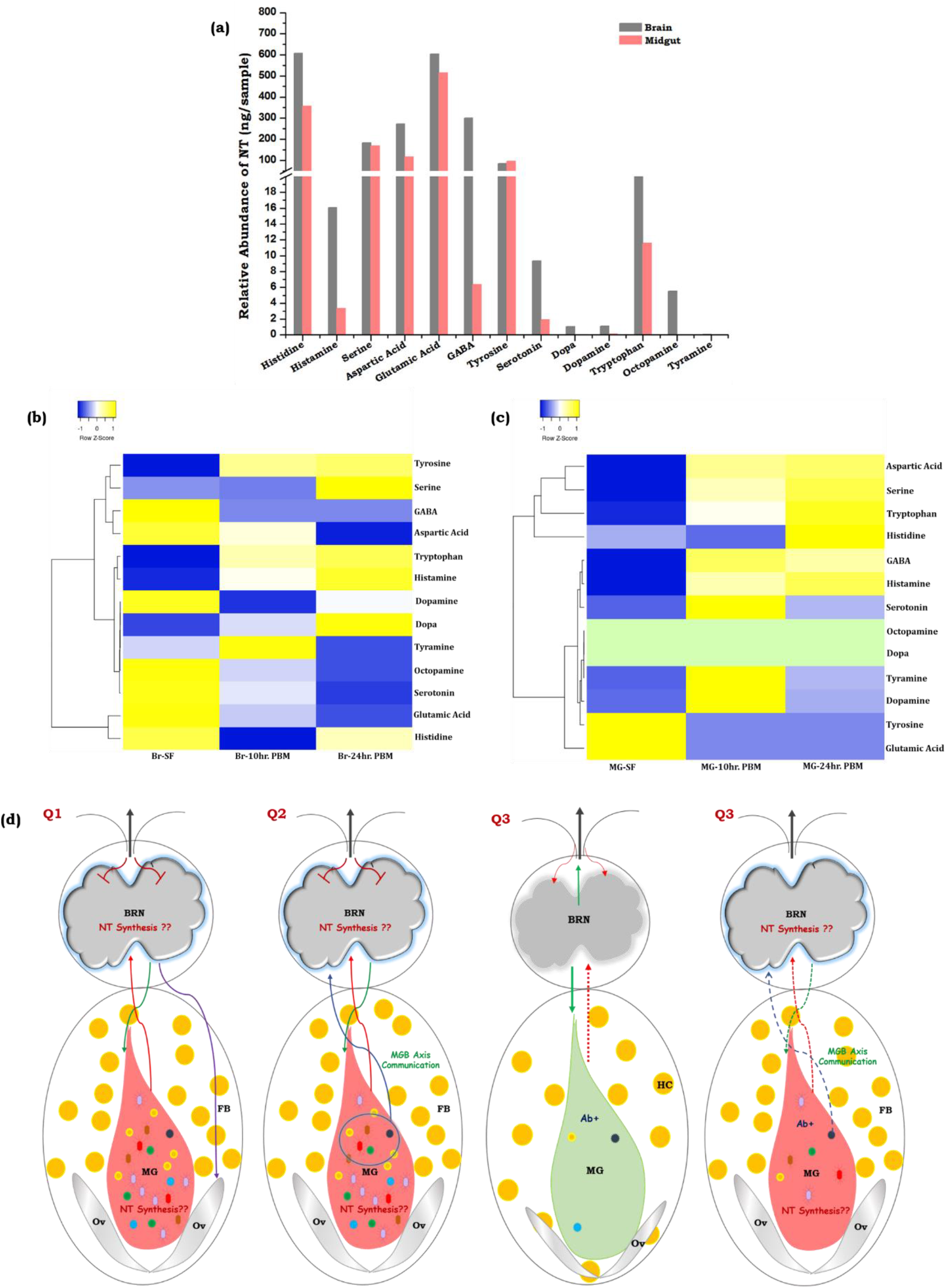
Gut-Brain-Axis (GBA) communication and neurotransmitter (NT) estimation in mosquito *An. culicifacies*. (a) Comparative analysis of NT abundance in the naïve mosquitoes’ brain and midgut; (b) Heatmap showing the alteration of neurotransmitters level in mosquito brain tissue. NT levels were measured by LC-MS from the brains of naïve (sugar-fed) and blood-fed females (10 and 24 h PBM) (n = 65, N = 2). Statistically significant differences in the amount of metabolites were tested by p-values (p ≤ 0.005) that are deduced by two-way ANOVA and Tukey’s test; (c) Heatmap of neurotransmitters levels of mosquito gut tissue that vary during the metabolic switch. NT levels were measured by LC-MS from the gut of naïve (sugar-fed) and blood-fed females (10 and 24 h PBM) (n = 50, N = 2). Statistically significant differences in the amount of metabolites were tested by p-values (p ≤ 0.005) that are deduced by two-way ANOVA and Tukey’s test. (n = number of mosquitoes from which the respective tissue was dissected and pooled for each independent experiment; N = number of biological replicates); (d) Pictorial presentation demonstrating GBA communication in response to gut-metabolic switch in mosquitoes. Q1: Blood-feeding pauses external stimulus-guided neuro-olfactory responses, but may shift brain engagement through the vagus pathway (red arrow) to regulate actions in the distant organs such as the midgut (green arrow) and ovary (purple arrow). Here, we questioned whether increased levels of amino acids in the gut during blood meal digestion may act like an NT. Q2: Do blood-meal-induced gut flora proliferation (different colored shapes indicate diverse microbial flora) influence GBA communication in mosquitoes. Q3: Whether gut-bacterial removal by antibiotic treatment confers the establishment of microbiome-gut-brain axis (MGB) communication. BRN: Brain, MG: midgut, FB: Fat body, Ov: Ovary, Ab+: Antibiotic positive/treated.

### Symbiotic gut flora influences gut-brain axis communication

The mechanism of gut-brain axis communication in vertebrates primarily involves neuronal stimulation through the vagus nerve, endosymbionts mediated regulation of the gut endocrine system, and other biochemical pathways[18, 19, 55]. Previous literature suggests that mosquito gut endosymbionts regulate many biological functions such as mosquito immunity, blood meal digestion, and ecological adaptation[49, 56]. Ingestion of protein-rich blood meal favors the rapid enrichment of gut microbiota[21], but whether it also affects the nexus of communication between the gut and brain remains elusive.

Therefore, to uncover the gut microbiome complexity and establish their possible relations with neurotransmitter abundancy, we evaluated the nature and diversity of gut microbiome population dynamics alteration in response to blood-feeding. A comparative metagenomic analysis unraveled that the naïve sugar-fed mosquito harbors 90% of the Enterobacteriaceae family of gram-negative gamma-proteobacteria such as *Enterobacter cloacae* complex sp., *Chonobacter sp*., *Escherichia coli*; 6% Psedomonodales family of gram-negative gamma-proteobacteria such as (a) *Acinetobacter sp*. members e.g. *Acinetobacter guillouiae*, *Acinetobacter iwoffii*, (b) *Pseudomonas aeruginosa sp.* group e.g. *Pseudomonas alcaligenes*, *P. nitroreducens*, *P. veronii*, *P. stutzeri* and *P. viridiflava*; and other bacterial family member of Bacteroidetes e.g. Flavobacteriacae - *Chryseobacterium sp*., *Elizabethkingia meningospetica*; beta-proteobacteria-Alcaligenaceae-*Alcaligenes faecalis* (Fig. 6/ Fig. S4a, b, c). However, surprisingly blood feeding not only supresses Enterobacteriaceae family member by 50%, but favors rapid proliferation of Pseudomonadales to 46% of the total community, where we observed dominant association of *Pesudomonas sp.*, *Acinetobacter johnsonii*; *Acinetobacter rhizosphaerae*, and other members from Alpha-proteobacteria family such as *Sphingobium sp.*, *Gluconacetobacter diazotrophicus*, *Achromobacter sp.*, *Sphingomonas azotifigens, Methylobacterium sp.* as well as Beta-proteobacteria-Burkholderiales family members such as *Acidovorex sp.*, *Delftia ramlibacter*; *Janthinobacterium lividum* (Fig. S4a, b, c). Our microbial profiling data further suggested that blood meal significantly alters the abundance of the gram-negative bacteria such as *Pseudomonas and Elizabethkingia* (Fig. 6), than gram-positive bacteria such as *Agromonas* and *Rubrobacter* (Actiobacteria) (Fig. S5).

**Fig. 6:**
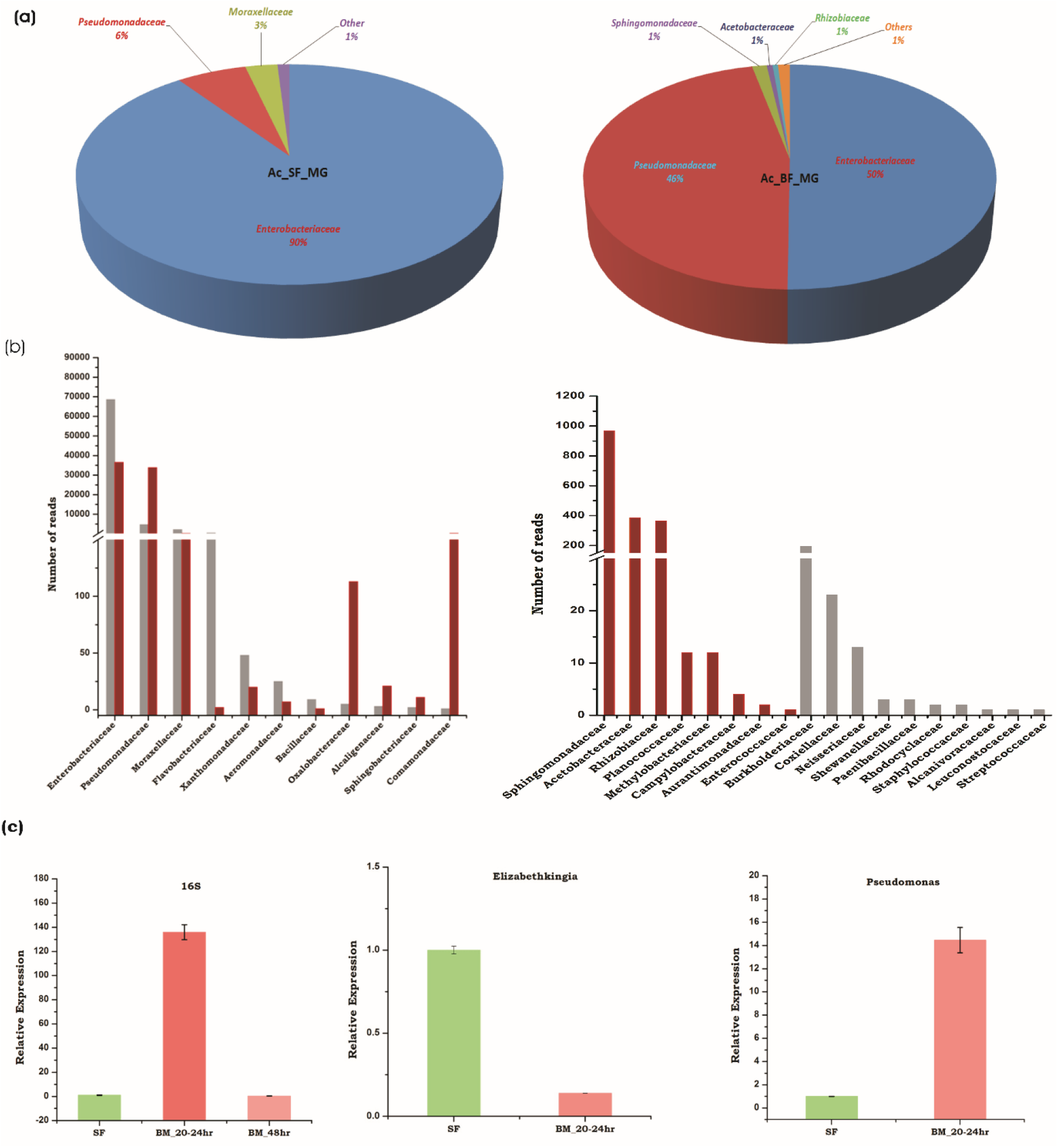
Comparison of gut-metagenomes in the naïve sugar-fed and blood-fed mosquito *Anopheles culicifacies*: (a) Pie charts showing the major bacterial families under the two feeding status (b) Number of reads based comparative bar graphs showing common and unique families microbes (c) Relative quantitative distribution of microbiota based on 16SrRNA based expression in the midgut of *An*. *culicifacies* in response to sugar and post blood feeding (20-24hrPBM, 48hr PBM); Relative abundance of *Elizabethkingia* and *Pseudomonas sp. bacteria* in sugar-fed and blood-fed (20-24hr PBM) condition.

Although the correlation of microbiome-gut-brain axis communication in the blood-feeding mosquitoes is yet not fully established, however, we opined that amino-acids resulting from rapid digestion of protein-rich blood meal, and its metabolite products may serve as an additional potent source of neuromodulators[57]. Here, our observation of Enterobacteriaceae family member abundancy and low NTs level in the gut than the brain of naïve sugar-fed mosquitoes indicate the basal-level of gut-brain-axis communication is enough to maintain physiological homeostasis. However, a rapid proliferation of Pseudomonadales family members, and a multi-fold enrichment of NTs in the gut, while mild suppression of the majority of NTs in the brain except for Histamine, Tyrosine and Tryptophan of the blood feed mosquitoes suggests that members of Pseudomonas species, may likely have a neuro-modulatory role in protein-rich diet-induced gut-brain-axis communication.

Furthermore, to test and evaluate the effect of gut flora removal on the neurotransmitters dynamics, we performed an absolute quantification of the potent neuroactive molecules and compared their levels in the gut and brain of the naïve and antibiotic-treated mosquitoes. A significant elevation of tryptophan and consequent downregulation of serotonin levels in both the gut and brain of aseptic non-blood fed mosquitoes (Fig. 7a, b), corroborate with the previous observations that depletion of microbial flora may significantly delimit the de-novo-synthesis of serotonin, resulting in increased tryptophan concentration in the gut and brain[58]. Additionally, we also observed that antibiotic treatment not only caused a notable increase in histidine and histamine levels in both the gut and brain, also favored an exclusive induction of Dopa, and significant enrichment of GABA in the gut of the aseptic mosquitoes (Fig. 7a, b). Together, these data indicated that gut bacteria removal may also influence the systemic level of amino acid concentration (Fig. 7a, b).

**Fig. 7:**
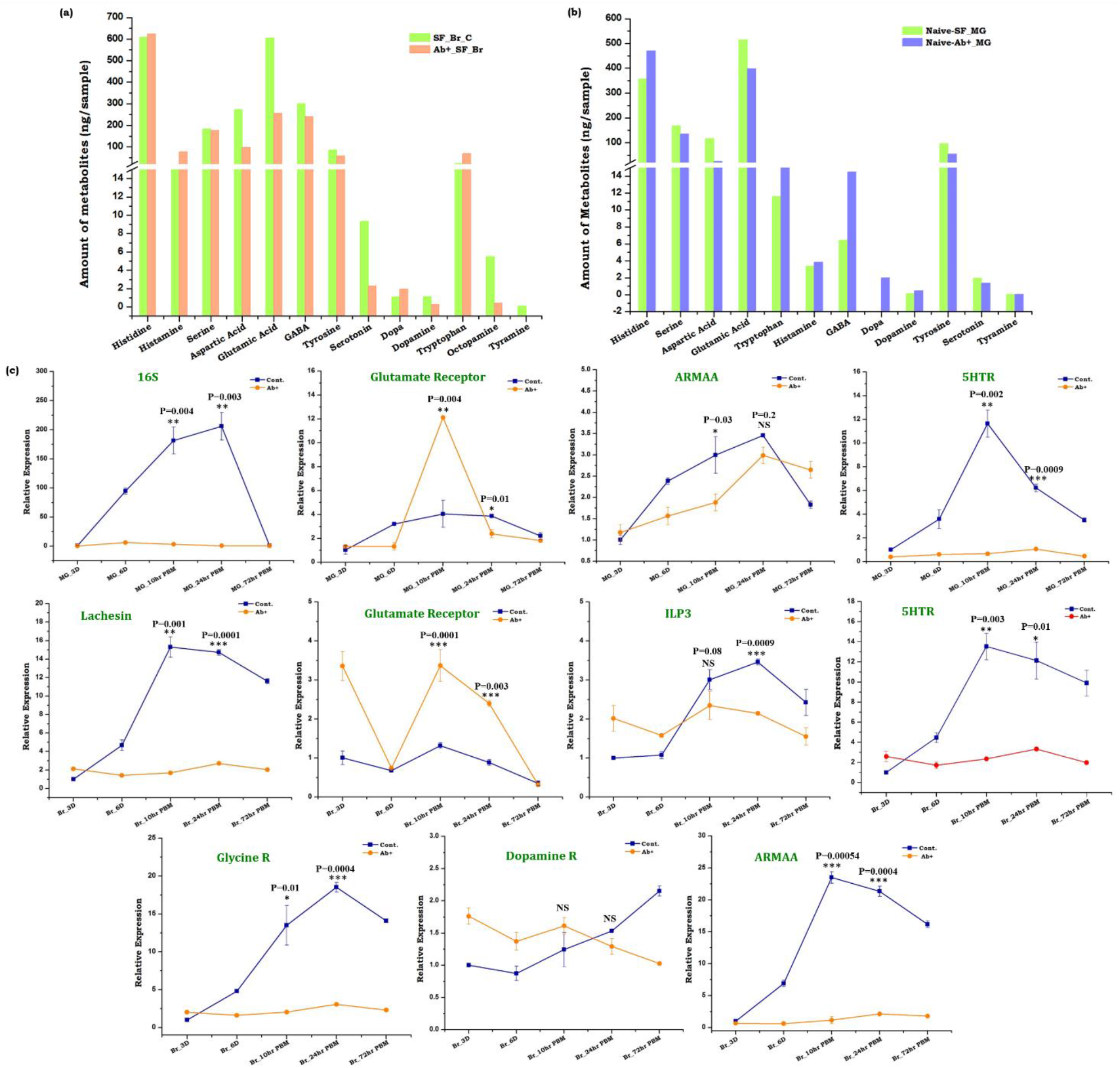
Establishing Microbiome-Gut-Brain-Axis (MGB) communication in mosquitoes. (a) Absolute quantification of the neurotransmitters (NT) in the brain tissues of naïve sugar-fed and antibiotic-treated mosquitoes (n = 65, N = 2); (b) Quantitative estimation of the neurotransmitters (NT) in the gut tissues of naïve sugar-fed and antibiotic-treated mosquitoes. Statistically significant differences in the amount of metabolites were tested by p-values (p ≤ 0.005) that are deduced by two-way ANOVA and Tukey’s test, (n = 50, N = 2); (c) Relative expression profiling of the 16S gene to show the population of microbial flora and other neuro-transcripts in the gut and brain of naïve and antibiotic-treated mosquitoes undergoes metabolic switch. Statistical significance of differences of the respective genes in control (without antibiotic) and aseptic mosquitoes (antibiotic-treated) were tested by the t-test. (n = number of mosquitoes from which the respective tissue was dissected and pooled for each independent experiment; N = number of biological replicates).

To test how blood-feeding influences gut-brain axis communication, again we quantified and compared the level of the neurotransmitters of naïve and antibiotic-treated blood-fed mosquitoes. A similar pattern of NTs synthesis was observed in both naïve blood-fed and antibiotic-treated blood-fed mosquitoes, but the level of modulation is heightened in antibiotic-treated blood-fed gut and brain (Fig. S7, Table S4). To further support the above observation, we also monitored and compared the expression patterns of neurotransmitter receptor genes (Glycine R, glutamate R, serotonin R, dopamine R), insulin-like-peptide, and one of the junction protein gene (lachesin) in the gut and brain of naïve vs. antibiotic-treated mosquitoes (Fig. 7c). Consistent with NT quantitative data, the respective receptor genes also showed a significant difference in their abundance throughout the gut-brain axis. We also noticed a differential expression pattern of ILP3, ARMAA (Aromatic-L-amino-acid decarboxylase/serotonin synthesizing enzyme), and lachesin transcript between naïve and antibiotic-treated mosquitoes undergoing metabolic switch event (Fig. 7c).

With our current data, we propose that a bi-directional gut-brain axis communication may exist to manage complex gut immune-physiological responses *via* gut-microbiome association during the blood meal digestion process in gravid females. Although, it is yet to clarify how this cross-talk directly influences brain-specific responses such as mood and cognition.

### The mosquito brain maintains basal immunity

The immune system plays a crucial role in maintaining brain health by protecting it from both external and internal stress[59]. Since the central nervous system and the immune system are the most energy-consuming organs, we consider that the immune system may play an important role to overcome blood-meal-induced metabolic stress, such as oxidative stress, osmotic stress, and elevated levels of dietary heme molecules. To trace the possible linkage of the brain-immune function, we identified and cataloged a total of 913 immune transcripts from brain tissue transcriptome data (Fig. 8a). Among the 18 classified immune family proteins, autophagy, CLIP-domain serine proteases, and peroxidases were observed the most predominant, accounting for more than 50% of the total immune transcripts. Furthermore, a comparative transcript abundance analysis showed that blood meal may cause a moderate change in the immune transcript expression (Fig. 8b). Increased percentage of peroxidases and CLIP-domain serine protease transcripts in the blood-fed brain suggested that these immune transcripts may prevent brain tissue from oxidative stress-induced damage and facilitate its recovery (Fig. 8b). Further, functional analysis of the immune transcripts in the central nervous system may unravel the novel regulatory mechanism of the immune genes to maintain the brain in shape.

**Fig. 8:**
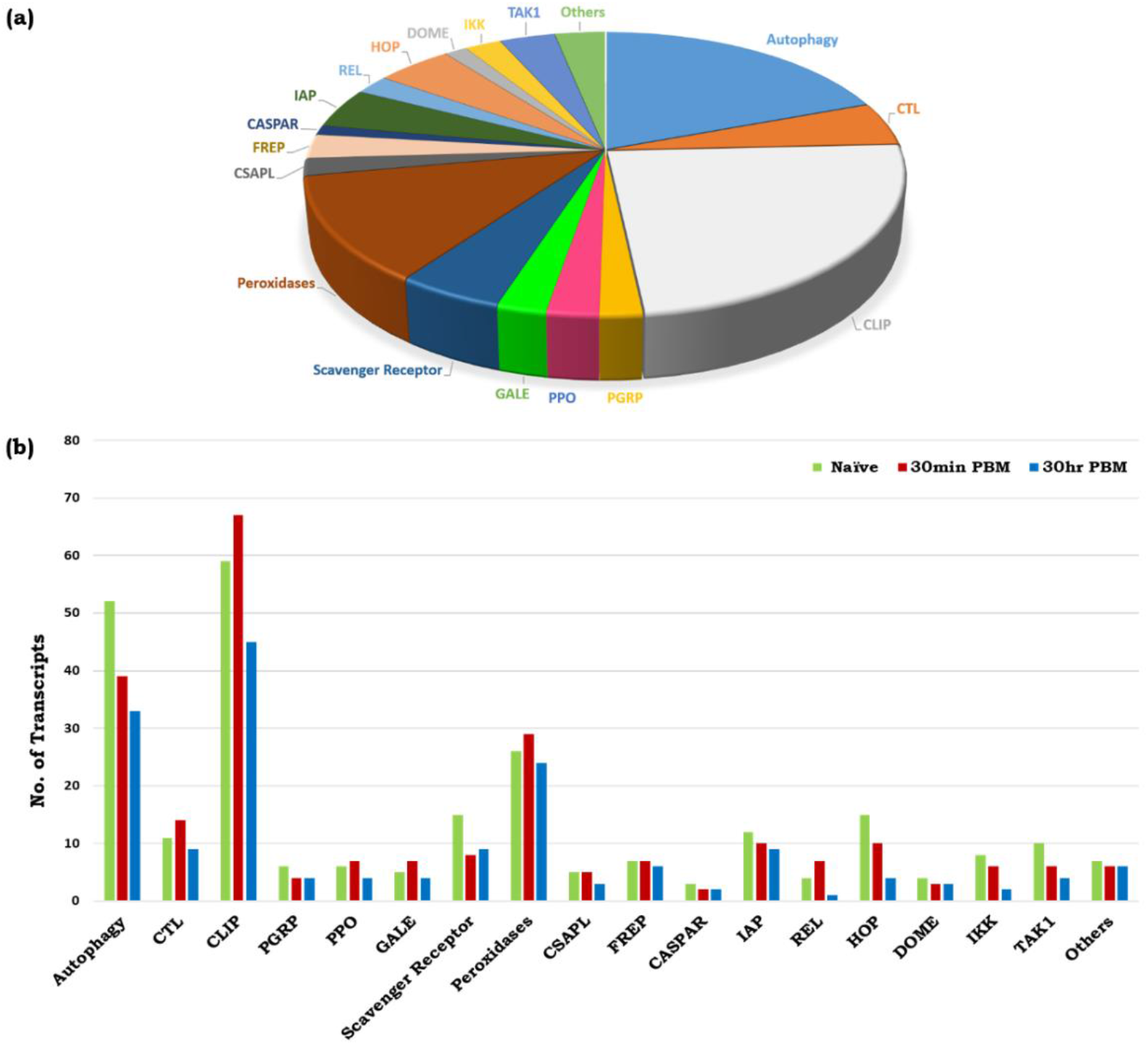
Molecular catalog of brain-specific immune transcripts. (a) Molecular catalog of the different classes of immune genes expressed in brain tissue; (b) Differential expression patterns of the brain immunome as determined by the number of sequences that appeared in each RNA-Seq data of naïve and blood-fed mosquito brains.

## Discussion

Host-seeking and blood-feeding behavior evolution make it difficult to resolve the complexity of decision-making neuro-actions in hematophagous insects[3]. Recently, *Benjamin J. Matthews et. al.*, cataloged hundreds of genes that are differentially expressed in the blood-fed brain[60], of which the brain-encoded neuropeptide Y has been suggested to play a crucial role in host-seeking suppression following blood feeding[4]. But, in-depth analysis of the gut-metabolic-switch-mediated modulation of brain function is unexplored. Our study attempts to establish a molecular relationship of gut-brain-axis (GBA) communication, and explore a possible functional correlation of gut-endosymbionts on neuro-transmitters dynamics influencing GBA communication.

### Gut-metabolic switch modulates the brain’s energy metabolism and functional engagement

To understand that how the engorgement of the gut with blood meal modulate brain functioning, we performed a comparative RNA-Seq analysis of naïve sugar-fed and blood-fed of adult female mosquitoes’ brain in *An. culicifacies*. In contrast to the pre-blood meal olfactory responses, which are significantly influenced by age/sex/circadian even in the absence of host-exposure[1, 29, 30], we did not observe any significant alteration of neuro-modulator genes expression in the non-blood-fed (host-unexposed) aging *An. culicifacies* mosquitoes. The onset of host-seeking behavior coincides with the age-dependent gradual change in the relative abundance of gene transcripts and increase in sensitivity of the olfactory sensory neurons (OSNs), where older females are likely to be more responsive to salient human odorants, and hence active host-seeker than teneral young female mosquitoes[61, 62]. Thus, our observation of a limited change in the neuro-modulatory genes expression in the mosquito *An. culicifacies*, suggests that cognitive learning and memory response towards host-attraction are poorly developed until mosquitoes are exposed to hosts.

Post-blood-feeding an exclusive induction of oxidation-reduction family proteins and a comparative pathway analysis predicts that blood meal may enhance the brain’s energy metabolic activities. Shreds of evidence from *Drosophila*, vertebrates, and limited studies in mosquitoes also suggest that altered metabolic physiology influences the cross-talk between the brain and peripheral tissues for the maintenance of systemic energy homeostasis[12, 35, 55, 63–65]. To perform this action, continuous neuronal stimulation is required, which consequently increases the energy demand of the brain[64, 66]. An enrichment of the fructose-mannose metabolic pathway and persistent elevation of PGC-1 and oxoglutarate dehydrogenase gene provides evidence of escalated energy metabolism and enhanced mitochondrial activity in blood-fed mosquitoes brain[31, 32]. In this context, it is plausible to propose that hyper mitochondrial activity may increase the ROS level which could have a deleterious impact on neuro action. Our observation of the unique appearance of the pentose phosphate pathway and glutathione peroxidase transcript (Oxidation-reduction category gene), along with the upregulation of CLIP-domain serine proteases and peroxidases immune transcripts may attribute to the scavenging of ROS generated due to enhanced mitochondrial activity. Moreover, blood-meal-induced expression of amino acid transporters and trehalose transporter indicated that both trehalose and amino acids may serve as a raw material for enhanced energy metabolism. Furthermore, we also observed a significant alteration of transcripts involved in intracellular signaling than neurotransmitter receptors in the blood-fed mosquitoes' brains. Together these data, we hypothesize that an internal nutritional stimulus may shift brain engagement from external communication to inter-organ management, which requires a rapid and continuous synaptic transmission, neurotransmitter recycling, and axo-dendritic transport, resulting in enhanced energy metabolism in the brain[64, 65].

### Neuromodulatory responses establish brain-distant organ communication

To support our hypothesis, we profiled a selected class of neuromodulators, neuropeptides, and neurohormones gene expression in the brain and correlated their impact on distant organs. Corresponding to the innate physiological status, we observed a time-dependent change in the expression pattern of the respective transcripts in the brain, and other targeted tissues of mosquito such as midgut, Malpighian tubule, and ovaries. But, in turn, how these neuromodulatory responses reinforce brain action remains unknown. Recent studies in *Drosophila* suggest that leukokinin neuropeptide regulates protein diet-induced post-prandial sleep and minimized movement[67]. We also observed a transient increase in leukokinin, and its receptor gene in the brain, and sustained up-regulation of the leukokinin receptor gene in the gut till 30h of blood-feeding. These data support the idea that until the blood meal gets digested in the gut, the brain may undergo ‘food coma’ and restrict external communication, but may actively engage in managing inter-organ communications (through ILPs and other neuro-hormones e.g. DH44, OEH, etc.). Compared to the brain, significant modulation of neuro-modulators, and sustained expression even in the gut of decapitated blood-fed mosquitoes, further suggested a specialized ability of the gut to serve as “second brain” possibly to share and minimize the function of the primary brain [16]. Taken together, we interpret that gut-metabolic-switching may favor the establishment of a bidirectional ‘gut-brain-axis’ communication in the gravid female mosquitoes, though the detailed molecular mechanism is yet to unravel.

### Neurotransmitter signaling and microbiome alteration influences Gut-brain-axis communications

Neurotransmitters, including both biogenic amines and amino acids, are well-known endogenous chemicals, that influence rapid inter-organ signal transmission and decision-making abilities[68, 69]. To clarify and establish a possible functional correlation between the gut metabolic switch and gut-brain axis communication, we quantified the levels of neurotransmitters secreted from both gut and brain tissues. When compared to the naïve sugar-fed status, an unusual and paramount shift from the brain to the gut was observed for almost all the neurotransmitter levels after blood feeding. A significant upregulation of aspartic acid, glutamic acid, histidine, and histamine levels in blood-fed mosquito gut and brain may be a consequence of the rapid degradation of protein-rich blood meal in the mosquito gut[70].

Although the effect of tyrosine enrichment in the brain is intriguing, however, an undetectable level of tyrosine in the gut supports previous findings that the scavenging of toxic tyrosine from the gut is essential for the safeguarding journey of blood-fed mosquitoes[71]. A substantial body of literature also suggests that the biogenic amines such as dopamine and serotonin are the critical regulators of feeding, host-seeking, and cognitive functions[72–76]. Thus, it would be worth testing whether an increase in the precursor molecules of dopamine i.e. tyrosine, in the blood-fed mosquito’s brain, improves the cognitive power of the mosquito’s host-seeking and blood-feeding behavioral activities. Likewise, an enrichment of tryptophan, a precursor of serotonin, may favor the minimization of the host-seeking behavioral activities of gravid females (Fig. 5b)[77]. Additionally, ~25-fold upregulation of GABAergic neurotransmission upon blood-feeding in the midgut highlights its possible function in the regulation of innate immune response by activating the autophagy due to gut flora expansion (Fig. 5c)[78, 79].

Appreciably, a recent term ‘psycobiotics’, which aims to examine the influential effect of the microbiome on the gut-brain axis communication, is common to vertebrate’s neurobiology, but insects’ communities are least attended[80, 81]. In vertebrates, studies suggested that mediators of the microbiota–gut–brain-axis communication are usually affected by microbial metabolism which includes short-chain fatty acids such as butyrate, neurotransmitters e.g. serotonin and γ-aminobutyric acid (GABA), hormones e.g. cortisol, and other immune system modulators e.g. quinolinic acid[82]. Further research on vertebrates and fruit flies indicated that gut microbiota influences several behavioral physiologies, including host metabolism, appetite, mood, sensory perception, and cognition[83–87]. Recent studies in flies also demonstrated that gut commensal bacteria and the composition of dietary amino acid supplements greatly influence in shaping behavioral responses such as food choice and olfactory guided foraging decisions[86, 88, 89]. However, studies on mosquitoes’ gut-symbionts are predominantly limited to their impact on parasite growth and their potentiality for para-transgenic approaches[90].

Our observation of a rapid proliferation of *Pseudomonas bacterial sp*. in the gut of blood-fed mosquitoes may likely due to increased consumption of dietary Tryptophan for the synthesis of serotonin, correlating a possible cause for the suppression of appetite (Fig. 6a, c & Fig. 5c)[91, 92]. Additionally, a significant reduction of the excitatory neurotransmitters Glutamic acid and Aspartic acid in the brain may help to restrict the foraging behavior in gravid females[93]. However, intriguingly a parallel thousand-fold increase in aspartic acid in the gut is whether a result of gut-microbial metabolism and/or any correlation with gut-neuro-endocrine regulation for egg development remains uncertain. Previous biochemical characterization of Locust’s vitellogenin protein showed that it carries high content of aspartic acid, glutamic acid, and leucine[94]. An independent *in-silico* amino-acid composition analysis of mosquito *An. culicifacies* vitellogenin protein (AEO51020.1) also revealed a high content of aspartic acid (6.2%), glutamic acid (6.7%), phenylalanine (7.6%), and serine (8.7%). Furthermore, previous literature indicated that disruption of gut-microbiota by antibiotic treatment not only reduces the anti-*Plasmodium* immunity but also hinder the egg development in the blood-feeding mosquitoes[21, 95]. Therefore, we correlate that blood-meal-induced gut-microbial metabolism and activation of the vitellogenesis process may sequester the majority of amino acids to nurture the eggs[52]. But, the remaining fraction of amino acids either serves as an energy reservoir in the fat body [52] or functions as a neurotransmitter, possibly to maintain gut-brain-axis communication, though further studies are needed to prove these presumptions.

A noteworthy modulation of gut neurotransmitters reinforces us to test how the blood-meal induced rapid proliferation of gut flora also influence gut-brain axis communications. We disrupted the gut symbionts by providing an antibiotic diet supplement to the newly emerged mosquitoes for 4-5 days and observed a significant difference in the abundance of neurotransmitters in both the gut and the brain. Surprisingly, we also noticed that aseptic adult female mosquitoes carry assertive feeding behavior towards a vertebrate host (personal observation). In animals, an earlier study showed that germ-free mice exhibited stress-induced altered behavioral response, which was restored after complete microbiota recolonization[96]. Studies further signify that the microbial antigens such as lipopolysaccharide (LPS) and lipoteichoic acids generated in response to antibiotic treatment elicit immune responses and favors early development of the gut-brain axis communication via gut neuronal sensing [97]. In the mosquito *An. stephensi,* the antibiotic treatment also enhances the transcriptional responses of gut-immune peptides, but the effect on neuro-sensing remains unclarified[98]. We interpret that a higher abundance of histamine in the brain and GABA in the gut of antibiotic-treated mosquitoes may be accountable for the enhanced host-seeking behavioral activities, either directly through neuro-stimulation or indirectly through the vagal pathway[99, 100]. Furthermore, blood-feeding to aseptic mosquitoes resulted in a multi-fold up-regulation of serine and glutamic acid suggesting a limited usage of the respective amino acids, in the lack of a microbial population (Fig. S7) (Table S4), which consumes crucial amino acids to synthesize the building blocks of bacterial cell wall components in the healthy blood-fed mosquitoes[101, 102].

### Conclusion

The current investigation provides a novel insight that how gut-metabolic-switch-induced transcriptional modulation shifts mosquito’s brain engagement from external communication (pre-blood meal host-seeking and host selection) to manage inter-organ communication (post-blood meal physiological homeostasis) for the fitness of the mosquitoes. To the best of our knowledge, our data provide initial evidence that correlates the potential role of gut endosymbionts in microbiome-gut-brain-axis communication. We believe our conceptual framework may be valuable to modify mosquitoes’ olfactory perception and cognition through the alteration of gut bacteria, and hence for new vector control tool development.

## Data deposition

The sequencing data were deposited to National Centre for Biotechnology Information (NCBI) Sequence Reads Archive (SRA) system (BioProject accessions: PRJNA555826; BioSample accessions: SRR9853884 for Ac-Br-SF, SRR9853885 for Ac-Br-30Min, and SRR9853883 for Ac-Br-30hrs).

## Authors contribution statement

TDD, PS, KCP, YH, RKD: Conceived and designed the experiments: TDD, PS, ST, DS, VS, CR, SK, ST, JR; RD; contributed to design and performing the experiments, data acquisition, writing and editing; TDD, PS, YH, KCP, RKD: data analysis and interpretation, data presentation, contributed reagents/ materials/analysis tools, wrote, reviewed, edited, and finalized MS. All authors read and approved the final manuscript.

## Acknowledgment

We thank insectary staff members for mosquito rearing. We also thank Kunwarjeet Singh for technical assistance in the laboratory. Finally, we are thankful to Xceleris Genomics, Ahmedabad, India for generating NGS sequencing data and DNAXperts, Noida, Utter Pardesh for metagenomic data generation and analysis. We thank CCAMP Metabolomics Facility for performing neurotransmitter (NT) analysis. We are grateful to Padma Ramakrishnan, Technology Scientist, Mass Spectrometry Facility - Centre for Cellular and Molecular Platforms, Bengaluru for her inputs in the manuscript by providing the details of LC/MS-based NT analysis and also thank her for troubleshooting during our study.

