## Supplemental data for "Microbiome-Gut-Brain-Axis communication influences metabolic switch in the mosquito *Anopheles culicifacies*"

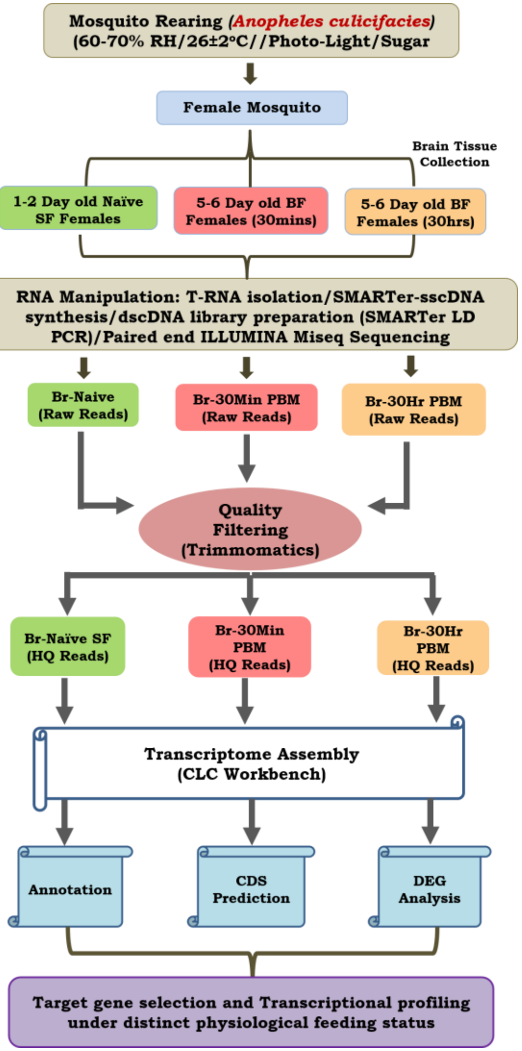


**Fig. S1:** A technical overview to decode the hard-wired genetic structure of brain system of *Anopheles culicifacies* mosquito.

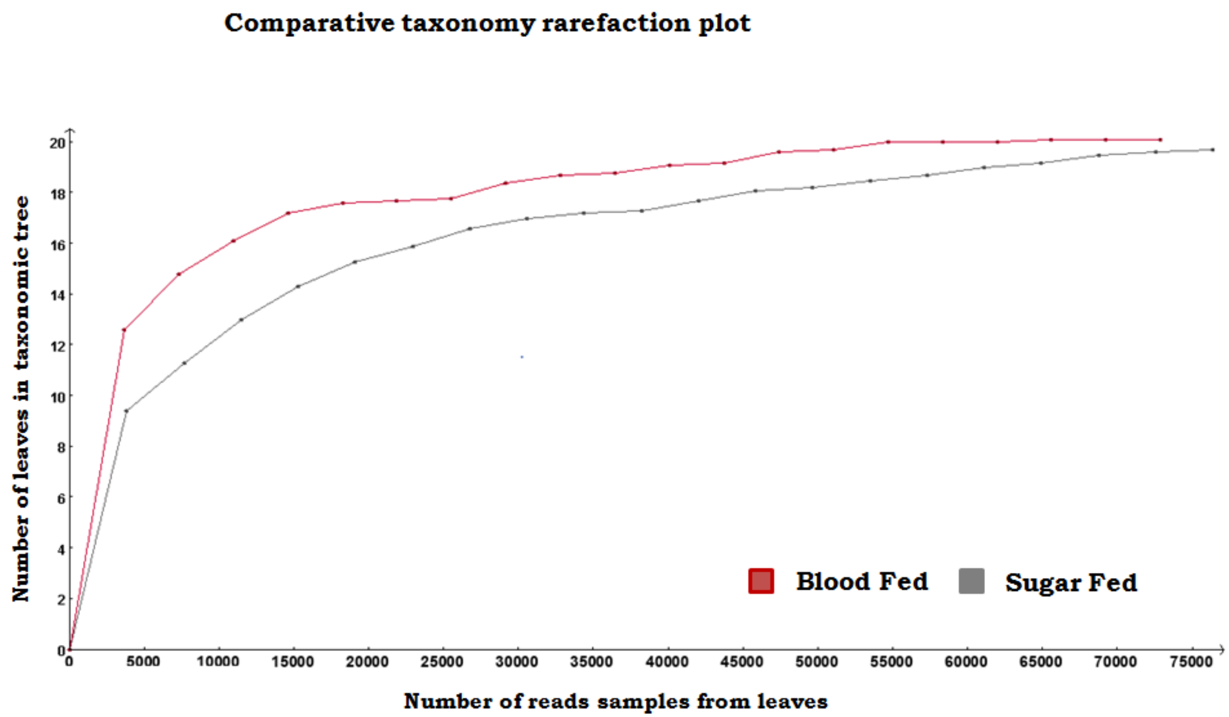


**Figure S2**: Rarefaction curve plot showing alpha diversity for the sugar fed (Gray line) and blood fed (Red line) at family level

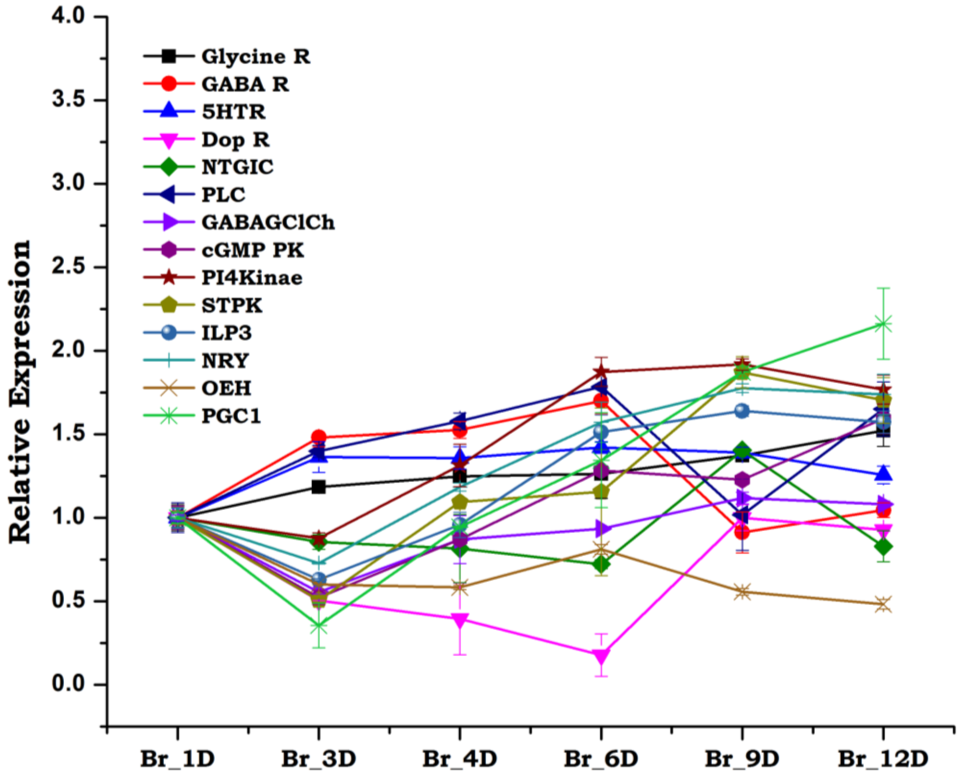


**Fig. S3:** Differential gene expression analysis in aging non-blood fed female mosquito brain. No significant modulation of neuronal genes was observed when compared to 0-1-day old teneral mosquitoes with 12 day-old non-blood fed mosquitoes. Statistical analysis using two-way ANOVA followed by Tukey Test has implied at 0.05 level, the expression pattern of the respective genes was not statistically significant in aging mosquitoes *(n = 25, N = 3)*.

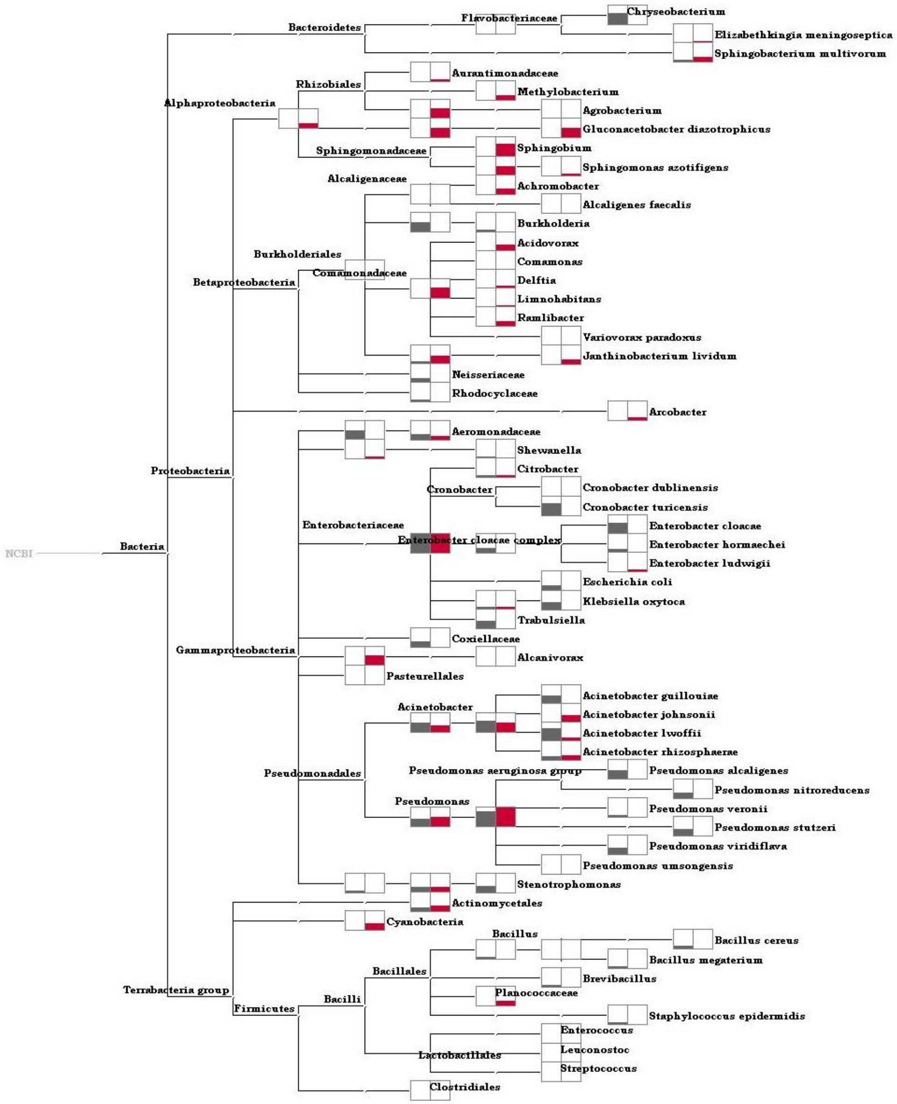


**Figure S4a:** Heat map of the comparative sugar fed (Gray bar) and blood fed (Red bar) midgut metagenomic data showing values at log scale for species level

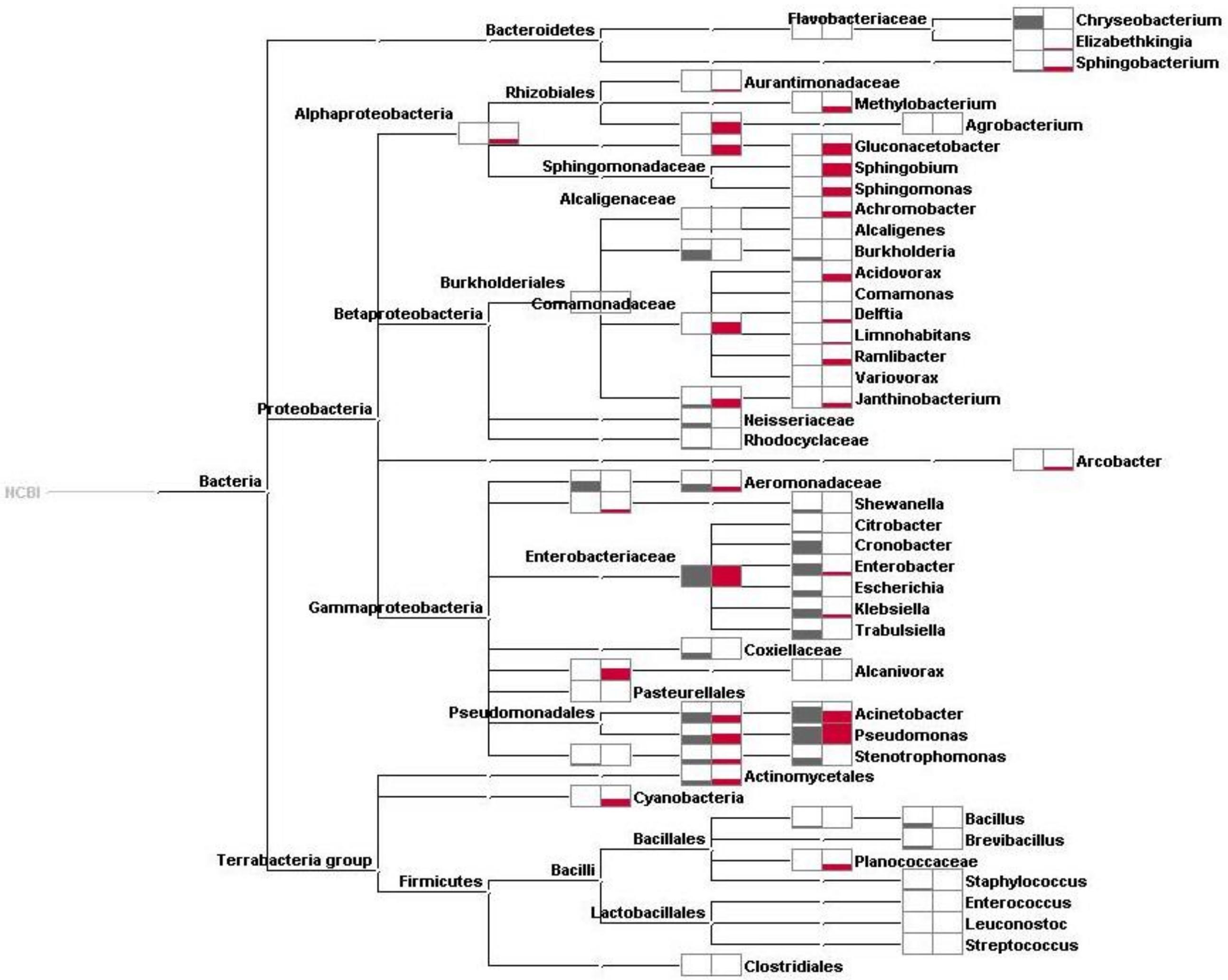


**Figure S4b:** Heat map of the comparative sugar fed (Gray bar) and blood fed (Red bar) midgut metagenomic data showing values at log scale for genus level

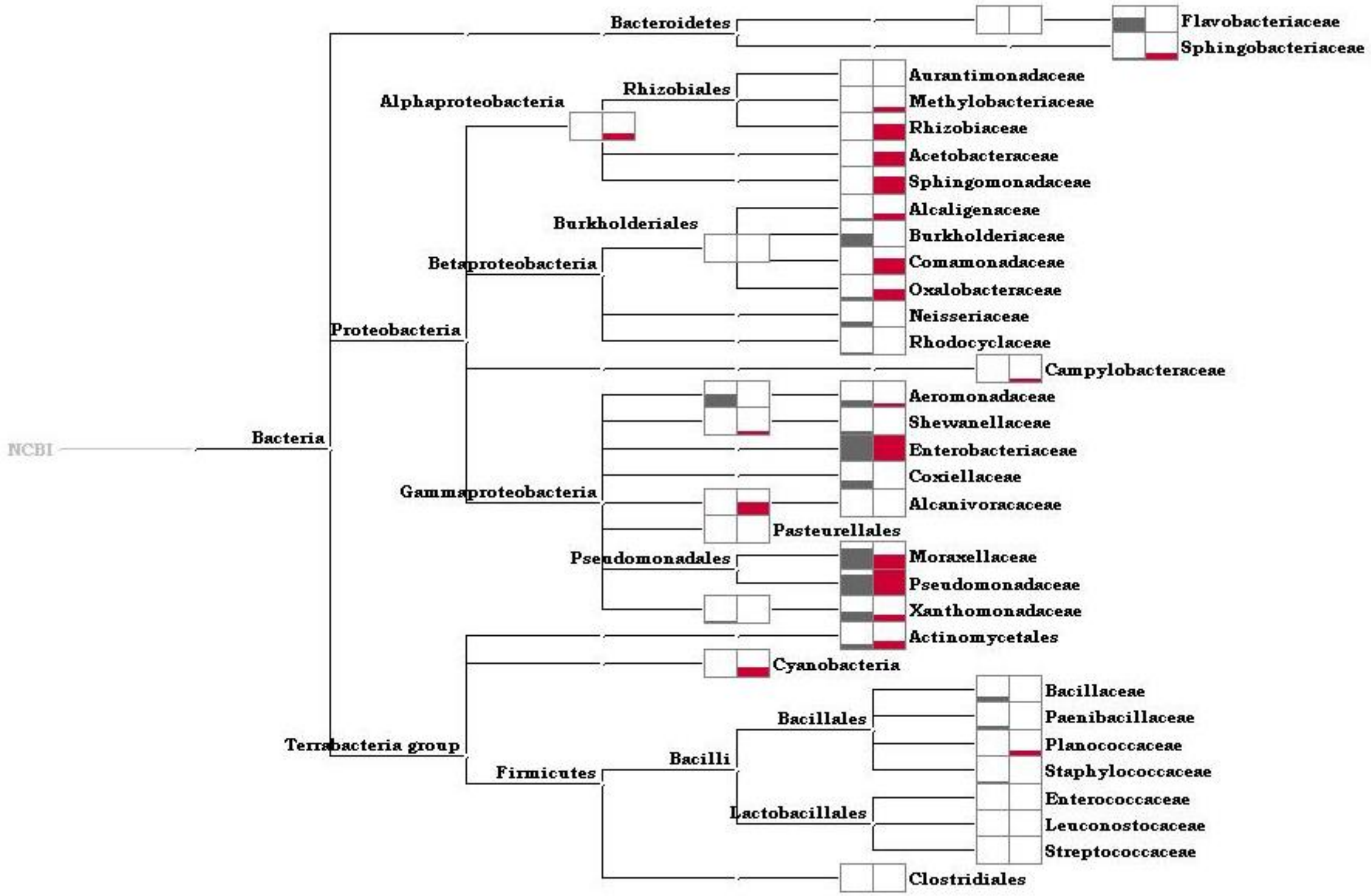


**Figure S4c:** Heat map of the comparative sugar fed (Gray bar) and blood fed (Red bar) midgut metagenomic data showing values at log scale for family level

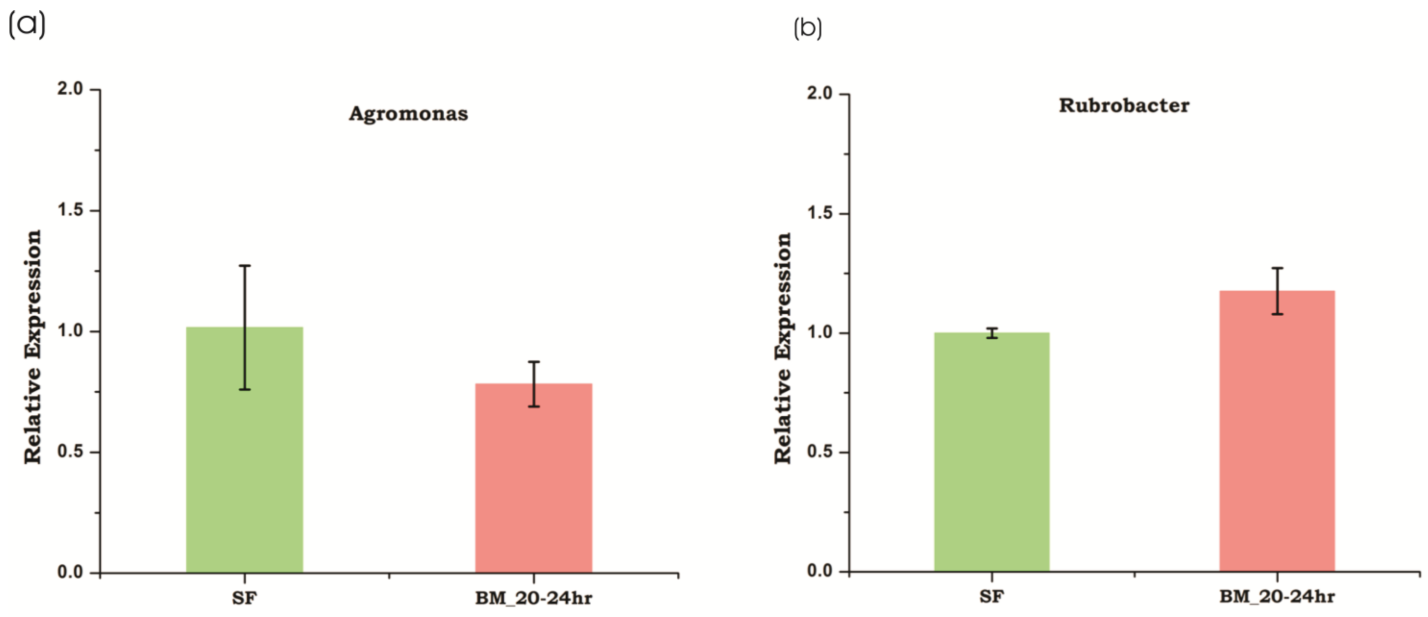


**Figure S5:** Relative expression of *Agromonas* and *Rubrobacter* (gram positive bacteria) at Sugar fed and 20-24hr Post blood fed condition shows no significant change during the two feeding status

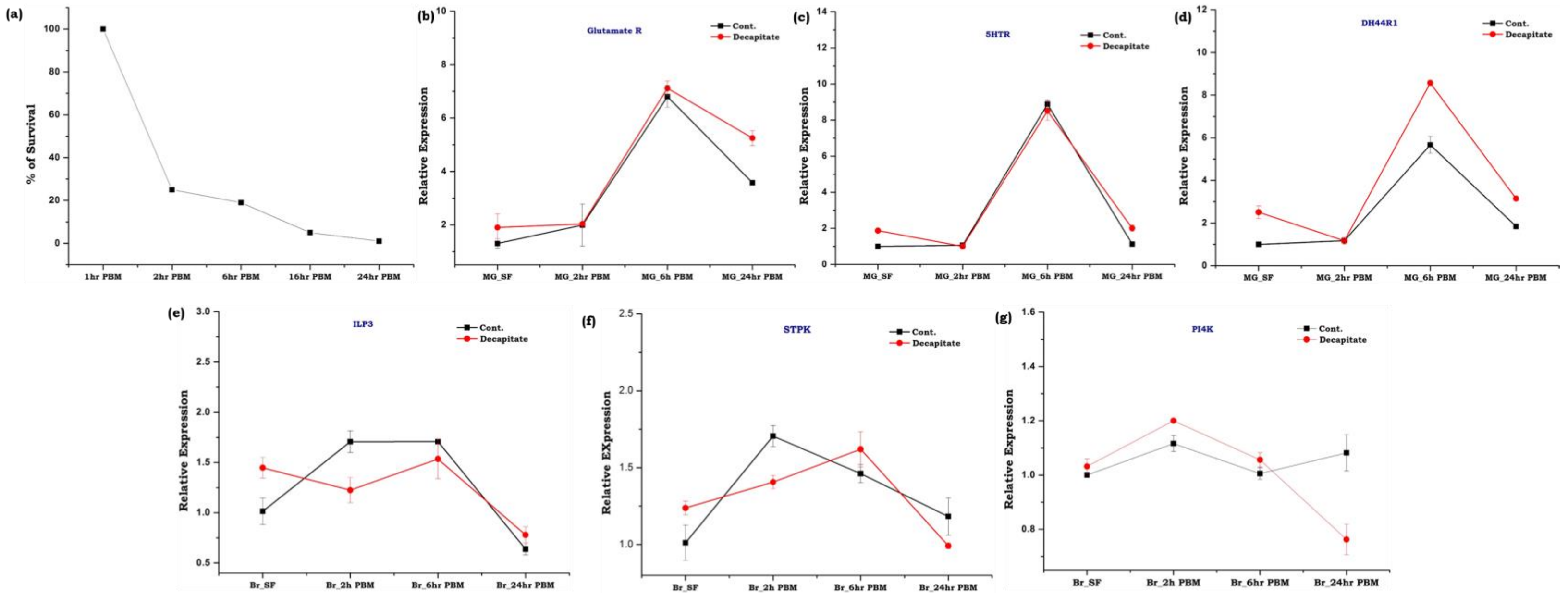


**Fig. S6: Transcriptional Response of neuromodulator receptor genes in naïve and decapitated blood-fed female *An. culicifacies* mosquitoes.** (a) Percentage of mosquitoes that survived till 24 h after decapitation which was performed after 1h of blood-feeding. 5-6 days old sugar-fed mosquitoes were provided blood meal and mosquitoes heads were decapitated after 1 h. from the full-fed gravid females. After that, the decapitated mosquitoes were kept in a cage for recovery and count the live (mosquitoes that vibrate/move their legs or other body parts are considered as live) and dead mosquitoes (non-movable mosquitoes with visible shrinkage of the body parts at the respective time points are considered as dead) at different time points until we observed 100% mortality. The percentage of survival was calculated until 24h after blood feeding. (b-d) Relative gene expression analysis of neuromodulator receptor genes in the gut of blood-fed and decapitated female mosquitoes. (e-g) Relative gene expression analysis of neuromodulator receptor genes in the gut of blood-fed and decapitated female mosquitoes. Statistical analysis using two-way ANOVA implied that at 0.05 level the expression level the respective genes in control and decapitated females are not statistically different.

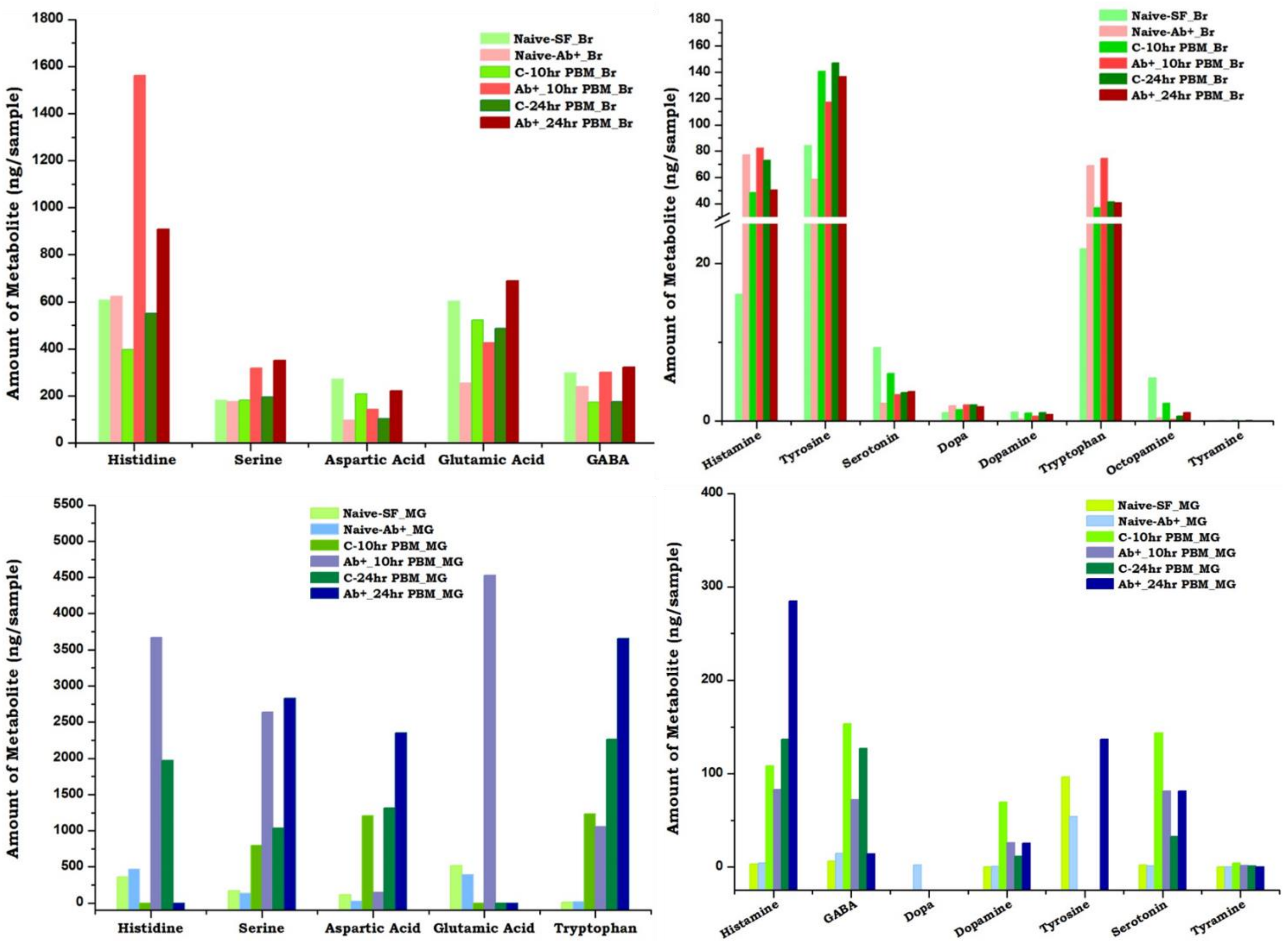


**Fig. S7:** Neurotransmitters dynamics of naïve and aseptic mosquitoes collected from sugar fed and blood fed conditions

**Table S1: List of Primer Sequence used in the study**

| Sl. No. | Primer Name and Sequence |
|---|---|
| **1.** | Actin_Fw: 5' TGCGTGACATCAAGGAGAAG 3'<br>Actin_Rev: 5' GATTCCATACCCAGGAACGA 3' |
| **2.** | Ac_PyruvateKinase_Fw: 5’ CGCACTTGATCTCCAAGTAT 3’<br><br>Ac_PyruvateKinase_Rev: 5’ TTCCAGCCAGTAACAACAA 3’ |
| **3.** | Ac_Solute Carrier 7_Fw: 5’ TCAATAGCTCCGAAATCAGT 3’<br><br>Ac_Solute Carrier 7_Rev: 5’ TGATAACGAACAGCAAGACA 3’ |
| **4.** | Ac_AATransporter_Fw: 5’ CAATGCCTATGGTTACAGGT 3’<br><br>Ac_AATransporter_Rev: 5’ GCTGGTAAGTGTCCTTCTTG 3’ |
| **5.** | Ac_TrehalaseTransporter_Fw: 5’ CGATGGGACTGTACTTCTTC 3’<br><br>Ac_TrahalaseTransporter_Rev: 5’ GTCTAGATCGGCGAAAAAC 3’ |
| **6.** | Ac_PGC1_Fw: 5’ ACCTTACGGTAAAATCGTCA 3’<br><br>Ac_PGC1_Rev: 5’ GTACGGTAGCTGATGTTCGT 3’ |
| **7.** | Ac_OxoglutarateDHS_Fw: 5’ GCAACTACTTCCATCTGCTC 3’<br><br>Ac_OxoglutarateDHS_Rev: 5’ GAGCCTTCAACAAGTCGTAA 3’ |
| **8.** | cGMP PK_Fw: 5’ GCGTTTGATTATCTGCACTC 3’<br><br>cGMP PK_Rev: 5’ AAGGACTCCAAGTGACCAGT 3’ |

| | |
|---|---|
| **9.** | GlutamateR_Fw: 5’AGTGGTATCAACGCAGAGTG 3’<br><br>GlutamateR_Rev: 5’ GAGTTTAAGCACTGCTCCAC 3’ |
| **10.** | Glycine R_Fw: 5’ GATACTGCCACTACCTCGTC 3’<br><br>Glycine R_Rev: 5’ CTTGGAGACCGAATTGAATC 3’ |
| **11.** | GABA R_Fw: 5’ CAGAACGAAGAAGGCTACTC 3’<br><br>GABA R_Fw: 5’ AGTATCCACGCATACTCAGC 3’ |
| **12.** | ARMAA_Decarboxylase_Fw: 5’GGTAACCAAGTCCTTCAGTG 3’<br><br>ARMAA_Decarboxylase_Rev: 5’ TAGAACAGACGACCTCGAAC 3’ |
| **13.** | ILP1_Fw: TCCACTACATGGAAAACTCC<br><br>ILP1_Rev: GTCATCAGTGCCTGGTAGAT |
| **14.** | ILP3_Fw: TAGCAATGATGAGTGGATGA<br><br>ILP3_Rev: ACAACACTCCTCTACGATGC |
| **15.** | Leukokinin_Fw: AAACATCGCATAGCAGAGAT<br><br>Leukokinin_rev: TCAGATAATCCTGCACCATA |
| **16.** | NRY_Fw: TACTGTACGGCTGGTTGAAT<br><br>NRY_Rev: TTAGTTCCGGCAGTGTTC |
| **17.** | OEH_Fw: GACAAGAATGCGGTGATAAT<br><br>OEH_Rev: CGTTGCTGTAGTAATCGAAG |
| **18.** | DH44R1_Fw: CTCGAAATAGAATGCTCCTG<br><br>DH44R1_Rev: AGATGACGATGAGGTAGGTG |

| | |
|---|---|
| **19.** | LKR_Fw: AAGAGGGAACACGACAAAC<br><br>LKR_Rev: GCTCGATATAATTGGTGGTC |
| **20.** | DH44_Fw: AACGAACAGGAAGATCTCAA<br><br>DH44_Rev: ATACCGTAGACGTACCGTGA |
| **21.** | CCHAR2_Fw: CCACTCCGAAAACTACAGAC<br><br>CCHAR2_Rev: GTGGCAGGAAGTAGTAAACG |
| **22.** | V-Type ATPase_Fw: TTACATGTACACCGATTTGG<br><br>V-Type ATPase_Rev: GACTTCATCAGACGTGACAG |
| **23.** | DopR_Fw:5’ GTTATGGGCGTGTTTATTGT 3’<br><br>DopR_Rev:5’ GCTGGTACTTGCGTCTTATC 3’ |
| **24.** | AKT Kinase_Fw: GATGGAGGAGGTAAAGTTCC<br><br>AKT Kinase_Rev: GAACTCACGGTCGAAGTAAC |
| **25.** | CYP314A1_Fw: GAGATTGCGCAAGAATTTAG<br><br>CYP314A1_Rev: GGAAGTTGTCCTCACTCTGA |
| **26.** | PTTH_Fw: CTTCACCTCTGAATTGCTTC<br><br>PTTH_Rev: ACAAGAAGACGGGTACTGTG |
| **27.** | KDNaCaExchanger_Fw: GTGAGATGGGTATCAGCAAC<br><br>KDNaCaExchanger_Rev: CTTCCAATCAAGTTTGAAGC |

| | |
|---|---|
| **28.** | GABA ClCh_Fw: 5’ GGAAGGTGTTTGGTAAGTCA 3’<br><br>GABA ClCh_Rev: 5’ GGTGATCGTGTTCGAGTAAT 3’ |
| **29.** | PLC_Fw: 5’ TGGATTCGTCCAACTATCAT 3’<br><br>PLC_Rev: 5’ TTCACGATCACCTCGTTC 3’ |
| **30.** | PI-4Kinase_Fw: 5’ ACATCATCTCCTCACTGTCC 3’<br><br>PI-4Kinase_Rev: 5’GTGTGCCACTGTTGTAATCA 3’ |
| **31.** | ST ProteinKinase_Fw: 5’TTTATAGTGCCGTGTGTTGA 3’<br><br>ST ProteinKinase_Rev: 5’CTTAATGTGGAACCGATCAT 3’ |
| **32.** | Trehalase_Fw: 5’ GAAGAGGACAAACAGGACTA 3’<br><br>Trehalase_Rev: 5’ GTTCCGGTAACCATAGAAC 3’ |
| **33.** | 5-HT Receptor_Fw: 5’ ATGATCTCGCGTAACTCCTC 3’<br><br>5-HT Receptor_Rev: 5’ ATCGGATTGACCAGACTGC 3’ |
| **34.** | TOR_Fw: GTAGAATGTTGGTGGTCGAT<br><br>TOR_Rev: ACCATCTGCTAGGTTATTGC |
| **35.** | Octopamine Receptor_Fw: 5’CTACTGGCGGATCTATCGGG 3’<br><br>Octopamine Receptor_Rev: 5’ TGGTGGAAGGCTGTGTTTTG 3’ |
| **36.** | Calcitonin R_Fw: AATAGAATGCTGGATGAACG<br><br>Calcitonin R_Rev: GGACGAAACGGTGTAAGTAT |

| | |
|---|---|
| **37.** | NTGated IonCh_Fw: 5’ ACGTTTCGAAAGTCAAACAC 3’<br><br>NTGated IonCh_Rev: 5’ GCTGTAGAATGCACAAATGA 3’ |

**Table S2: Comparative alpha diversity indices estimation of gut-bacterial population of naïve sugar fed and blood fed mosquito *An. culicifacies***

| **Sample** | **Taxonomy Rank: Class** | |
|---|---|---|
| | **Shannon index** | **Simpson index** |
| **Ac_SF_MG** | 0.088 | 1.019 |
| **Ac_BF_MG** | 0.225 | 1.063 |

**Table S3a: Annotation kinetics of RNA-Seq data**

| Molecular Features | Ac-Br-Naive | Ac-Br-30M PBM | Ac-Br-30Hr PBM |
|---|---|---|---|
| **Total No. of Raw Reads** | 5268211 | 3947521 | 3760078 |
| **Total No. of Contigs** | 32118 | 32984 | 38512 |
| **Total Transcripts** | 9460 | 9146 | 7387 |
| **Total BLASTx hits (NR)** | 8,668 (~91%) | 8,336 (~91%) | 6,548 (~88%) |
| **Transcripts with GO Match** | | | |
| **Molecular Function** | 4773 | 4556 | 3575 |
| **Biological process** | 4446 | 4299 | 3381 |
| **Cellular component** | 2523 | 2424 | 1888 |

**Table S3b: Percentage of differentially expressed transcripts**

| Sample | No. of Transcripts | Transcripts showing Differential gene Expression (DGE) | Upregulated Transcripts | Downregulated Transcripts | Percentage of Transcripts showing DGE |
|---|---|---|---|---|---|
| Ac_Br_naive vs Ac_Br_30min | (9460 + 9146) = 18606 | Total - 4747<br>Significant – 3183<br>Not significant - 1564 | 622 (3%) | 2110 (11%) | 14% CDS show differential expression |
| Ac_Br_Naive vs Ac_Br_30hr | (9460 + 7387) = 16847 | Total -3966<br>Significant – 3174<br>Not significant - 792 | 482 (2%) | 2469 (14%) | 16 % show differential expression |

**Table S4: Quantitative estimation of 13 different neurotransmitters in the brain and the gut of mosquitoes under different physiological conditions.**

| Name of NT | Control SF_Br | Ab+_SF BR | Control SF_MG | Ab+_SF _MG | 10hr PBM_Br | Ab+_10hr PBM_Br | 10hr PBM_MG | Ab+_10hr PBM_MG | 24hr PBM_Br | Ab+_24hr PBM_Br | 24hr PBM_MG | Ab+_24hr PBM_MG |
|---|---|---|---|---|---|---|---|---|---|---|---|---|
| **Histidine** | 607.77 | 622.52 | 356.52 | 469.70 | 397.58 | 1562.75 | DC | 3672.30 | 550.28 | 908.02 | 1968.73 | DC |
| **Serine** | 182.75 | 176.07 | 167.62 | 134.24 | 182.04 | 318.04 | 798.51 | 2636.08 | 197.18 | 350.75 | 1034.20 | 2830.31 |
| **Histamine** | 16.08 | 76.89 | 3.35 | 3.83 | 48.28 | 82.21 | 108.12 | 83.16 | 73.12 | 50.42 | 136.69 | 284.97 |
| **Aspartic Acid** | 272.35 | 96.74 | 115.69 | 24.02 | 209.89 | 143.05 | 1205.17 | 146.59 | 102.76 | 222.80 | 1312.08 | 2348.14 |
| **Glutamic Acid** | 604.78 | 255.68 | 513.97 | 397.82 | 523.25 | 425.24 | DC | 4530.44 | 486.92 | 690.13 | DC | DC |
| **GABA** | 300.01 | 240.89 | 6.38 | 14.50 | 174.40 | 300.76 | 153.31 | 72.01 | 175.86 | 321.81 | 126.79 | 13.94 |
| **Dopa** | 1.06 | 1.94 | DC | 2.00 | 1.45 | 2.04 | DC | DC | 2.08 | 1.84 | DC | DC |
| **Octopamine** | 5.49 | 0.39 | BLQ | BLQ | 2.29 | 0.20 | NF | DC | 0.65 | 1.11 | NF | NF |
| **Tyrosine** | 84.26 | 58.49 | 96.41 | 53.93 | 140.86 | 117.51 | DC | DC | 147.28 | 136.63 | DC | 136.63 |
| **Dopamine** | 1.12 | 0.28 | 0.13 | 0.46 | 1.00 | 0.62 | 69.17 | 26.02 | 1.06 | 0.83 | 11.59 | 25.18 |
| **Serotonin** | 9.33 | 2.25 | 1.92 | 1.35 | 6.06 | 3.39 | 143.62 | 81.14 | 3.59 | 3.72 | 32.70 | 81.37 |
| **Tyramine** | 0.08 | BLQ | 0.05 | 0.06 | 0.10 | 0.05 | 4.06 | 1.73 | 0.07 | 0.05 | 0.90 | 0.22 |
| **Tryptophan** | 21.84 | 68.84 | 11.59 | 16.44 | 36.93 | 74.66 | 1230.41 | 1054.85 | 41.40 | 40.70 | 2263.10 | 3650.53 |

*** BLQ= Below Limit of Quantitation**
**NF= Not Found**
**DC= Detected but not calculated due to the highly suppressed Internal Standard signal**

#### Representative UHPLC-MS/SRM chromatogram of Blank:

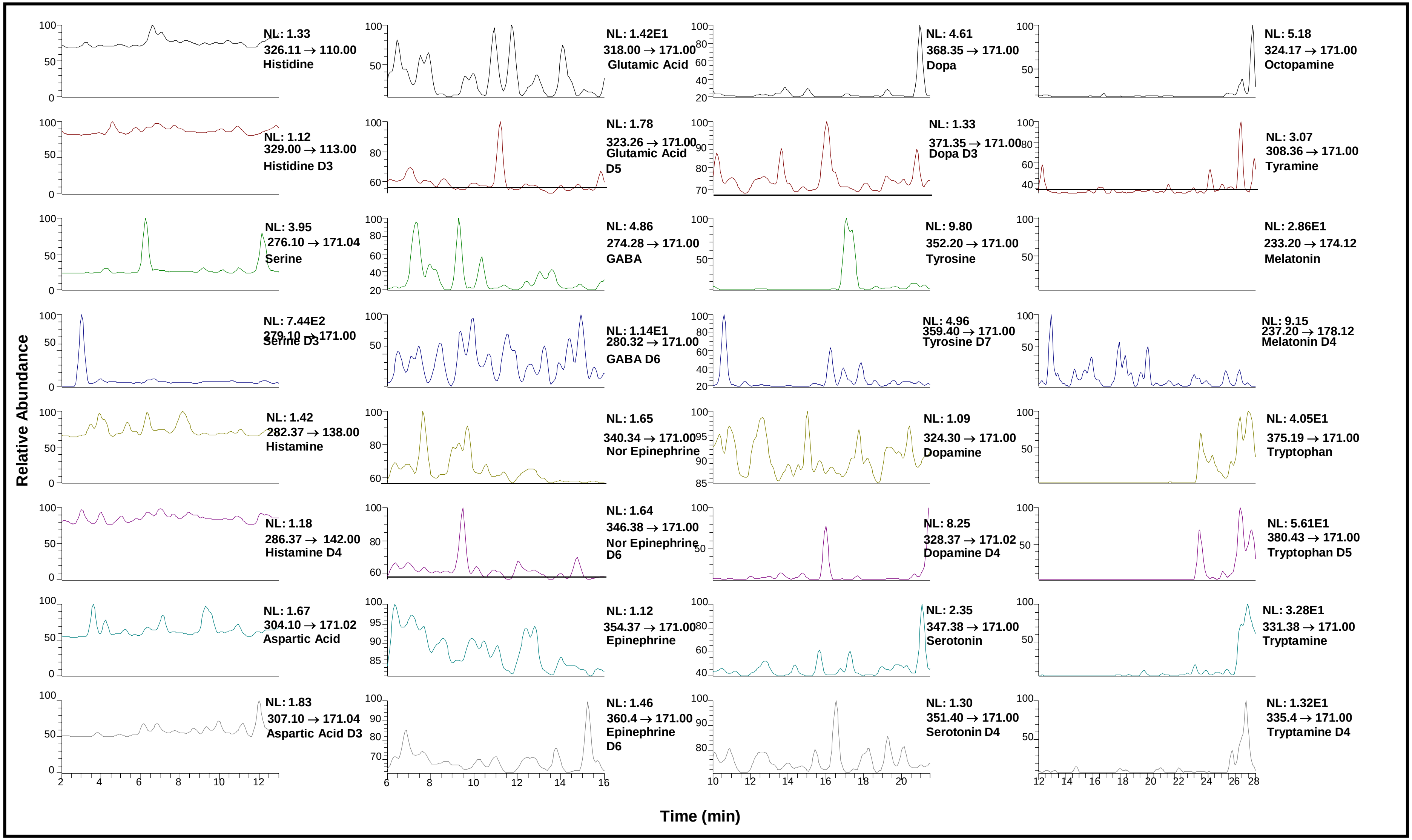

#### Representative UHPLC-MS/SRM chromatogram of Standard:

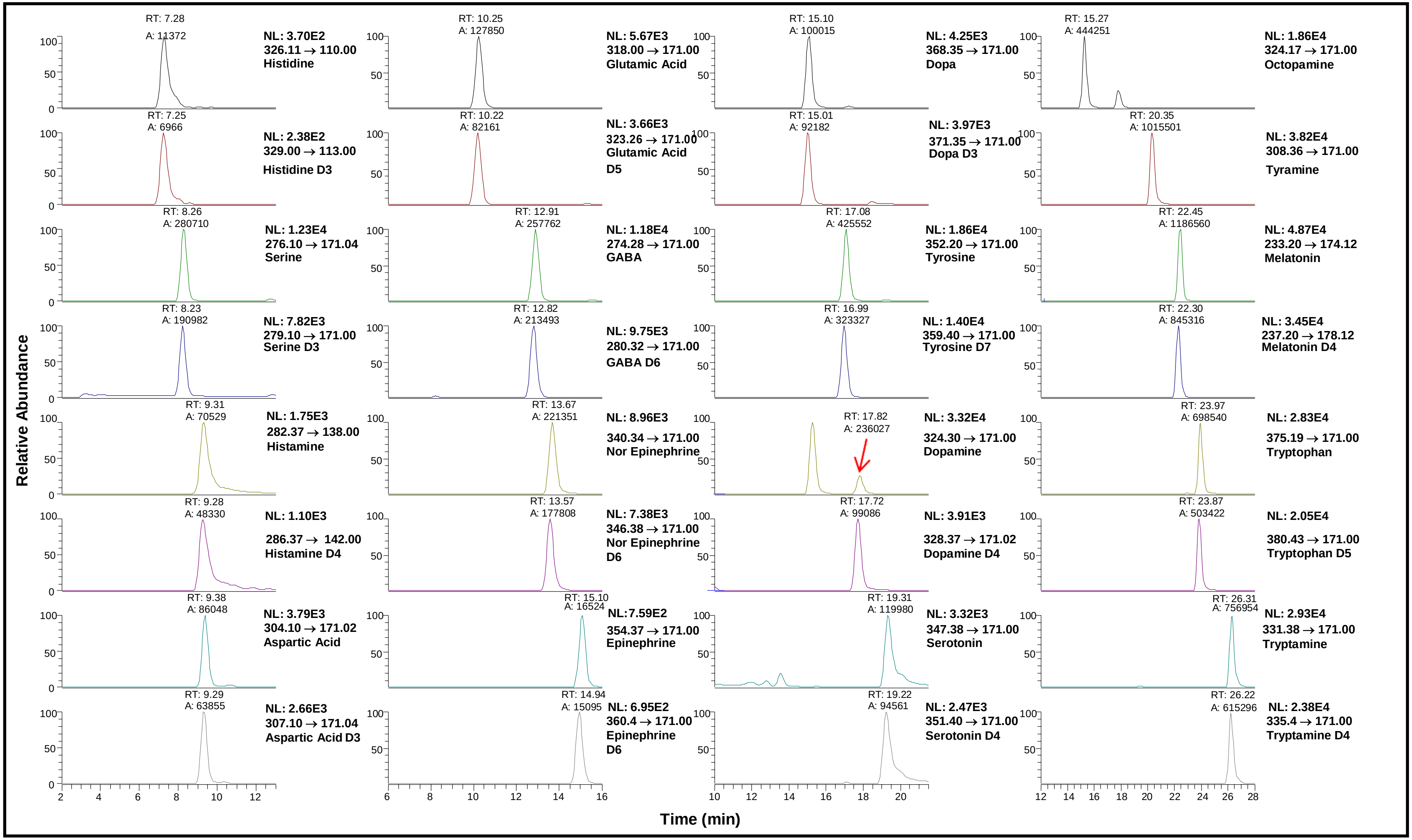

### Representative UHPLC-MS/SRM chromatogram of sample Ac_SF_Br:

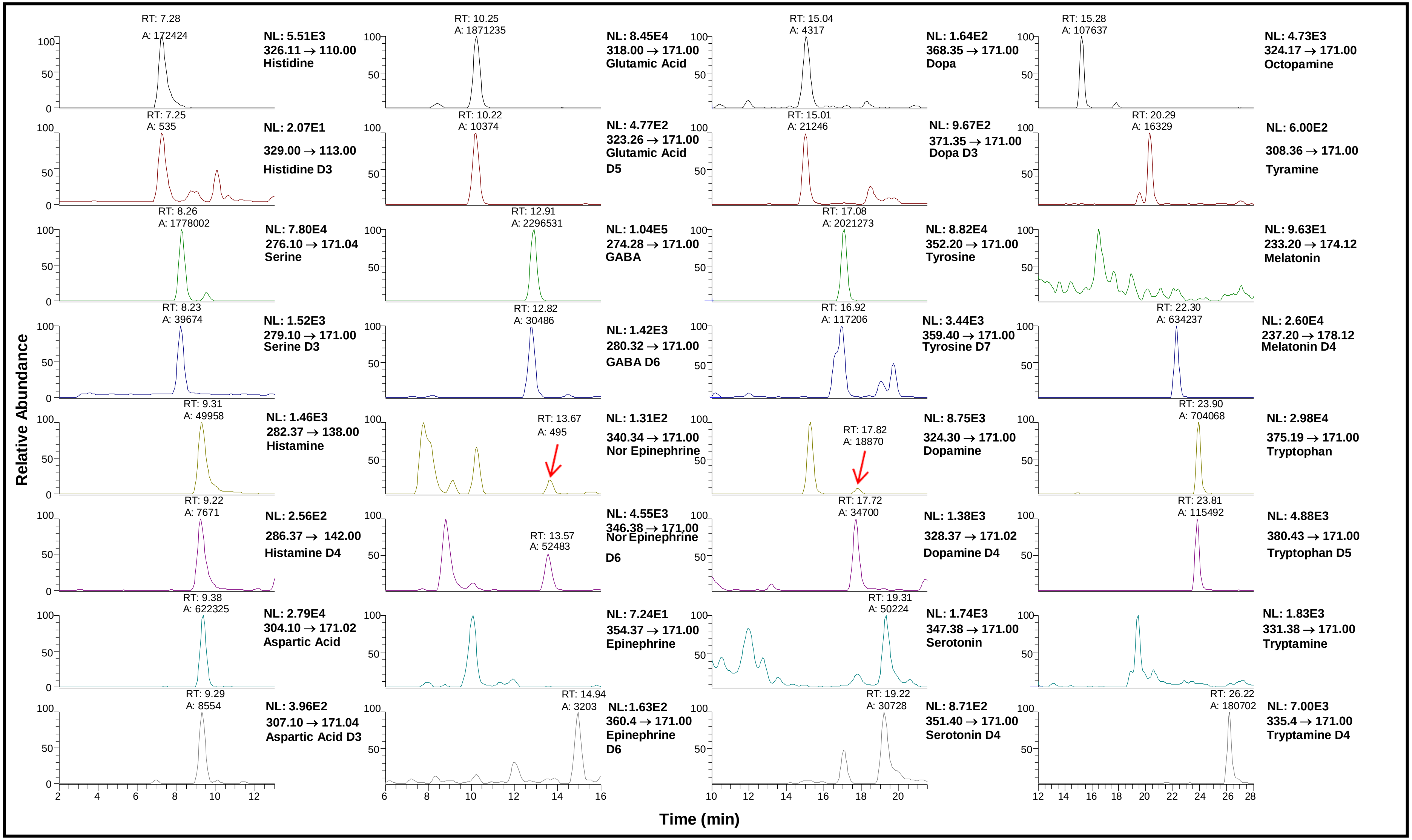

#### Representative UHPLC-MS/SRM chromatogram of sample Ac_SF_MG:

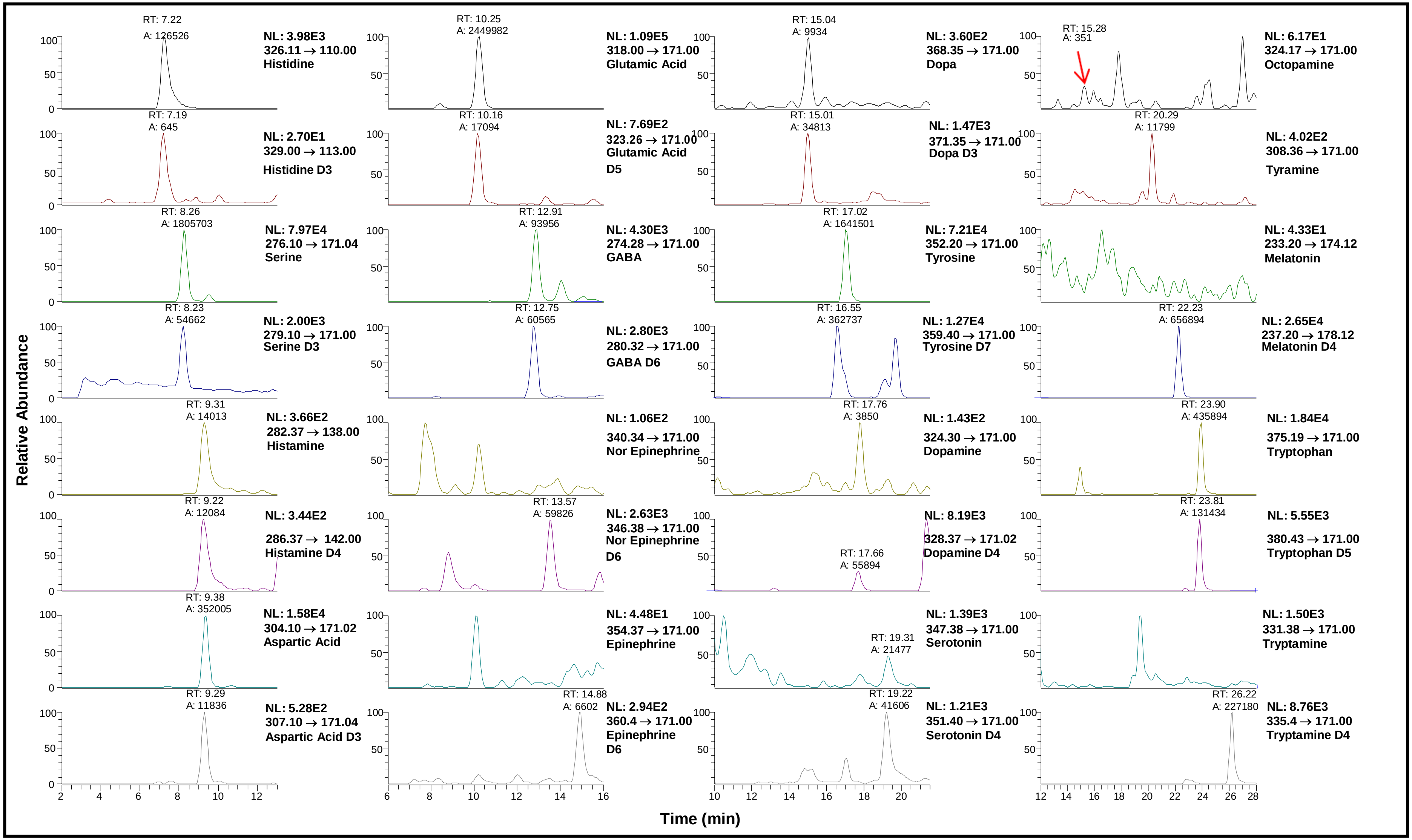
